# Extreme somatic mutation variation through time and space in walnut clones

**DOI:** 10.1101/2025.07.24.664184

**Authors:** Matthew W. Davis, Charles A. Leslie, Chaehee Lee, Evan Long, Li Meinhold, Megan Lorenc, Franklin Lewis, Patrick J. Brown, J. Grey Monroe

## Abstract

Many plants, unlike most animals, can reproduce as clones^1^. Cloning plants is critical for agriculture^2,3^ and biotechnology^4^, but the extent of somatic mutation arising during different propagation methods remains an important question for both fundamental research and agriculture. Here, we *discover* a surprisingly complex mutational history in a multi-decade natural experiment of isogenic walnut clones derived and maintained through three alternative methods: budwood propagation of field-grown trees, *in vitro* shoot culturing, and *in vitro* somatic embryogenesis. We generated a haplotype-phased reference genome assembly and revealed a >3500% increase in somatic embryo mutation rates compared to field-grown trees, along with a distinct mutation spectrum. The assembly also helped reveal extreme genomic instability in the somatic embryos, including multiple chromosomal duplications, megabase-scale deletions, telomere expansions, somatic recombination events, and ongoing transposable element activation. Our survey of somatic mutation also provides high-resolution insight into clonal stem cell dynamics, confirming the canonical meristem layers of flowering plants in the tree and shoot clones, while uncovering clear evidence of frequent single-cell bottlenecks in the somatic embryos. These discoveries inform practical questions about mutagenesis through plant tissue culture and serve as a benchmark to complement emerging paradigms of somatic mutation research in humans and other organisms.

## Introduction

Unlike most animals, many plants can be readily propagated as genetic clones^1^. Cloning has been a part of agricultural practices for thousands of years^2^, and millions of plants are now clonally propagated annually. Cloning facilitates the large-scale proliferation of preferred genotypes, but the extent to which genetic integrity is maintained in the face of somatic mutation is an open question. Clonal propagation is also critical for biotechnology, as it is an essential intermediate step in producing transgenic and gene-edited plants for many species^4–8^. Investigating somatic mutation in clonally propagated plants at whole-genome scales is critical for answering long-standing questions of fundamental and economic importance.

Plants can be cloned through various in-field and *in vitro* methods, including budwood cuttings, shoot culturing, and somatic embryogenesis^9^. Despite being clonally generated and assumed *a priori* to be genetically identical, plant clones occasionally display new and variable phenotypes, known as somaclonal variation^10,11^. This can result in poor field performance^12–14^, but it also may give rise to new crop varieties, as seen in wine grapes and oranges^15^. While some somaclonal variation is due to epigenetic differences^16^ or de-silencing of transposons^17^, much of it is likely due to *de novo* point mutations^18^ and other genomic changes. Here, we leverage new genomic sequencing technologies in walnut clones generated and maintained with different propagation techniques to compare mutation rates across time and space.

Somatic mutation in multicellular organisms creates natural mosaics of genetically distinct cell populations^19^. In mammals, somatic mosaicism has been investigated extensively to study cell lineage dynamics and development^20^, while similar investigations in plants represent a burgeoning frontier^21–23^. Mosaicism can present a significant challenge in plant biotechnology, as plants regenerated after transformation and gene editing are frequently chimeric^24^. To mitigate this, somatic embryos are often used because they are less prone to chimerism^25^. Though substantial effort has been focused on improving transformation and editing efficiency, the general genomic integrity of the clonal material warrants further attention. Incorporating somatic mosaicism as a framework for studying mutation in plants offers a powerful lens for uncovering fundamental plant biology and advancing methods in plant biotechnology.

To address these critical knowledge gaps, we combined short- and long-read sequencing to measure somatic mutation in a unique clonal walnut (*Juglans regia*) lineage established in the 1970s. These data show extremely elevated mutation and genomic instability in plant tissue culture, while simultaneously revealing clear developmental insights. This study represents the most exhaustive exploration of somatic mutation in plant tissue culture, capturing mutations from the single nucleotide level to whole-chromosome scales and providing a benchmark for assessing mutation in plant clones through time and space.

## Results & Discussion

### Mutation accumulation across clones

Since the 1970’s UC Davis has maintained clones of the world’s most popular walnut cultivar, ‘Chandler’. In 1995, a single clonal somatic embryo was generated from ‘Chandler’ anther tapetum and multiplied in culture by repetitive direct somatic embryogenesis to serve as a biotechnology resource^26^. Over the next three decades this culture underwent expansions and contractions, existing now as 12 subpopulations, all clonally derived from the single somatic embryo progenitor. This population provided us with a unique opportunity to investigate long-term somatic mutation.

Our initial goals were two-fold: to conduct a benchmark study of somatic mutation while addressing the long-standing question of how mutation impacts clonal plants. We began in 2021 by sequencing two field-grown trees with PacBio HiFi and two somatic embryos from the same subpopulation with Illumina Whole Genome Sequencing (WGS) (Fig. 1a, Supplementary Table 1).

**Fig. 1.**
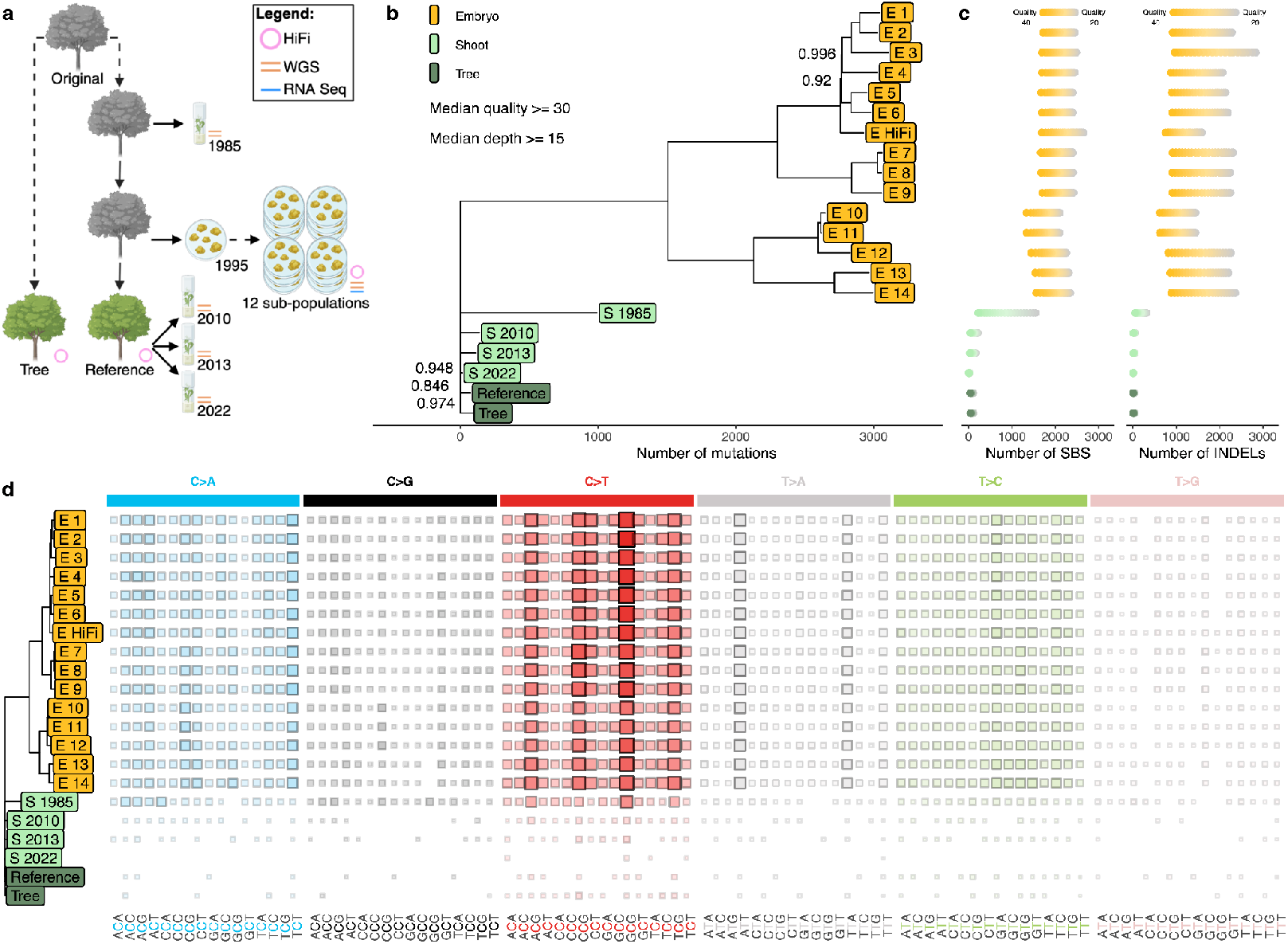
Somatic embryos exhibit high levels of mutation compared to shoots and trees. **a**, Schematic of the relationship amongst all clones. Icons denote trees, shoot cultures, and somatic embryos. Grey icons represent dead clones. Dashed lines indicate an unknown number of clonal propagations, solid lines indicate a single clonal propagation. **b**, Phylogeny of all sequenced clones constructed from *de novo* somatic SBS and InDels. Branch length is proportional to the number of mutations. Bootstrap support less than 1 is displayed. **c**, The number of SBS and InDels present in each sample at minimum median quality scores of 20-40, in increasing intervals of 1. **d**, The mutation spectra of SBS in the clones. Larger squares and darker color indicate higher occurrence of each context.

Combining HiFi and chromatin-capture Omni-C data, we constructed telomere-to-telomere haplotype-phased genome assembly for the walnut cultivar ‘Chandler’ from the same tree used to construct the previous reference genome assemblies^27,28^. The phasing accuracy of the assembly was confirmed using a genetic map (Supplementary Fig. 1-6, Supplementary Table 2-4). All *in vitro* clonal material was either introduced from this reference tree or its direct clonal progenitors (Fig. 1a) and are considered the same age, as they were derived from the same zygotic event.

Using a strict filtering strategy to remove ancestral heterozygosity, sequencing errors, and mapping artifacts, we identified a set of high-confidence *de novo* mutations (Methods, Supplementary Fig. 7, Supplementary Table 5). The two somatic embryos exhibited a surprising >3500% increase in mutation when compared to the trees. To confirm this unexpected result, we performed PacBio HiFi sequencing on a random somatic embryo from a random subpopulation, as well as Illumina WGS and RNA sequencing on a random somatic embryo from each of the 12 subpopulations. We also used Illumina WGS on clones generated with a different *in vitro* approach, sequencing shoot cultures introduced from nodal cuttings in 1985, 2010, 2013, and 2022 (Fig. 1a, Supplementary Table 2). Mutations in these samples were subsequently identified with the same pipeline.

We created a phylogeny of *de novo* mutations in all clones (Fig. 1b), which further confirmed the >3500% somatic mutation enrichment across embryo samples relative to the reference tree. This result held true regardless of sequencing method (i.e. short- vs long-read) or variant filtering parameters (Fig. 1b-c), and provides insight into the propagation history of the somatic embryos since their introduction three decades ago.

The elevated mutation rate in embryos could not be explained by relaxed purifying selection, despite fewer mutations observed within gene bodies compared to intergenic regions, which is consistent with patterns of neutral somatic mutation in other plants and greater DNA repair efficiency in gene bodies^29,30^ (Supplementary Fig. 8a). When compared to annual somatic and germline mutation rates in other plant species, the mutation rate in the somatic embryos was higher, though similar to the rate detected in rice callus^31–34^ (Supplementary Fig. 8b).

In the shoots, the number of mutations scaled linearly with time in culture (Supplementary Fig. 8c, Supplementary Table 5), with the longest maintained shoot culture (S 1985) showing a >1000% increase in mutation compared to the trees. This pattern of somatic mutation accumulation is comparable to the pattern seen in mammalian somatic tissues^35,36^.

The extreme differences in mutation accumulation were also reflected in the mutational spectra, as all somatic embryos exhibited a similar profile that was distinct from that of S 1985 (Fig. 1d, Supplementary Fig. 8d), despite somatic embryogenesis and shoot culture both being *in vitro* methods. The remaining shoot cultures and trees had too few mutations to individually assess differences in spectra (Fig. 1d, Supplementary Table 5).

Though all of these clonal samples are the same age, they exhibit vast differences in mutation rate. These results shed light on the mutagenic potential of tissue culture, with the mutation burden depending on cloning technique and time maintained in culture. The spectrum of mutation also appears to be dictated by the clonal propagation method, implying variability in DNA damage and repair, while suggesting developmental differences between the clones.

### Large-scale genomic instability

Chromosomal instabilities have been observed in cancer^37^, human somatic cells^38^, and various plant tissues^39–41^, and have the potential to impact the fidelity of clonal propagation in agriculture and biotechnology.

We assessed genomic instability with the sequencing depth of phased ancestral heterozygosity across clonal samples (Fig. 2a, Supplementary Fig. 9a). All somatic embryos carry a duplication of chromosome 9 haplotype B, and all but two carry a duplication of chromosome 4 haplotype A (Fig. 2b). We verified the most parsimonious explanation of the absent trisomy using allele frequency: the 4A duplication occurred once and was lost in the E 5 & E 6 clade, evidenced by the retention of 4A specific mutations in these individuals that occurred after chromosome duplication (Supplementary Fig. 9b-d, Supplementary Table 6). We dated the 4A and 9B trisomies using shared *de novo* mutations and discovered ~10-fold fewer mutations occurred before the duplication on 9B compared to 4A, indicating 9B duplicated much earlier in the somatic embryo’s history (Supplementary Fig. 9e, Supplementary Table 7). Additionally, there are incredibly few shared mutations on the duplicated 9B haplotype, suggesting that the duplication occurred either in the source material before the induction of embryogenesis or shortly after. Since the generation of this somatic embryo in 1995^26^, the walnut improvement program has been unsuccessful in initiating new ‘Chandler’ somatic embryo cultures despite numerous efforts, and the early onset of this duplication hints that it may be important for successful somatic embryogenesis in this genotype.

**Fig. 2.**
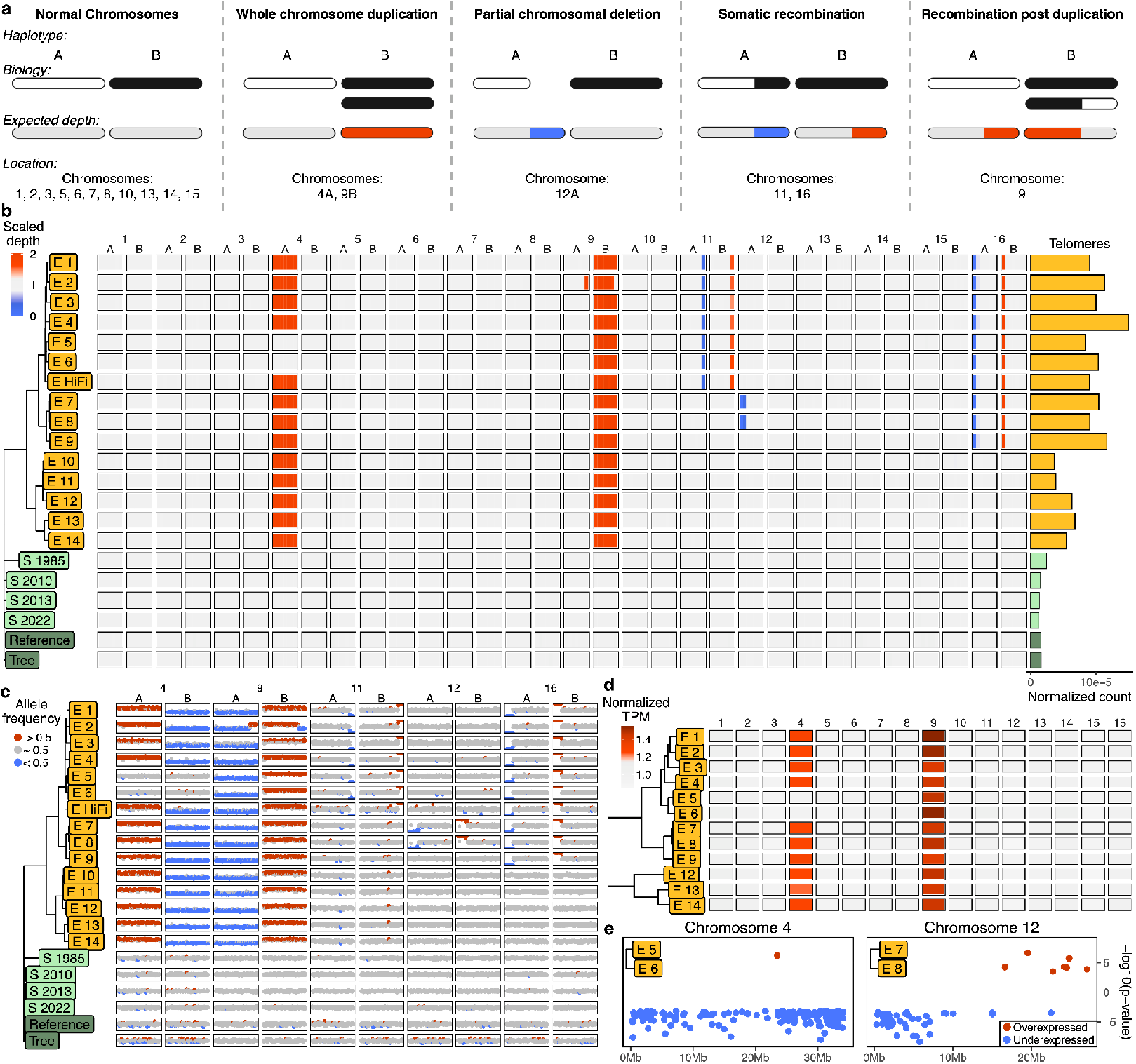
Genomic instability in the somatic embryo clones. **a,** Read depth schematic. **b**, Scaled read depth of ancestral heterozygosity in every chromosome of every clone. Red and blue represent high and low sequencing depth, respectively. Bars to the right display the number of telomeric repeats corrected by sequencing depth. **c**, Allele frequency of ancestral heterozygosity in chromosomes exhibiting genomic instability. Red and blue represent windows with mean allele frequency significantly greater or less than 0.5, respectively. **d**, Normalized expression of entire chromosomes in the somatic embryos. Red indicates higher than average expression. **e**, Comparison of individual gene expression across clades of embryos. Red and blue points are comparatively over- and under-expressed genes, respectively.

A partial chromosomal deletion of 12A, a somatic recombination between the chromosome 9 haplotypes in the trisomy, a somatic recombination between the chromosome 11 haplotypes, and a somatic recombination between the chromosome 16 haplotypes were also found in the somatic embryos (Fig. 2b, Supplementary Fig. 10a). Notably, there was a lack of genomic instability in the shoot cultures and trees.

We verified the instabilities using allele frequency and sequencing depth (Fig. 2c, Supplementary Fig. 10b), and further explored their effects on gene expression in the somatic embryos. The duplicated chromosomes exhibited higher gene expression relative to diploid chromosomes (Fig. 2d), and the clades of somatic embryos lacking the 4A duplication or with the 12A deletion exhibit lower expression. Considering these structural variants alone, we identified thousands of genes whose expression is likely affected by somatic mutation accumulation.

The somatic embryos also had more telomeric repeats than the shoot and tree clones (Fig. 2b, Supplementary Fig. 10c, Supplementary Table 8). Interestingly, one of the two major clades of somatic embryos had more telomeric repeats than the other clade (Supplementary Fig. 10d,e). S 1985 also had more telomeric repeats when compared to the other shoot cultures and trees, and telomere length in the shoots appears to correspond with time in culture (Supplementary Fig. 10f).

All instances of genomic instability in the somatic embryos are congruent with the SBS and InDel phylogeny, giving us additional confidence in our results and further insight into the clones’ shared history. These analyses also provide additional evidence that recombination occurs in plant somatic cells, a phenomenon that has been proposed to occur through the life cycle of a plant^42,43^, as a DNA repair response^42,44^, or as a pathogen stress response^45^. The lack of instability in the shoot cultures and trees further underscores the mutational consequences of specific clonal propagation methods.

Clone age has been associated with reproductive success^46^ and these genomic instabilities impact the expression of thousands of genes. We have attempted to germinate a few randomly selected somatic embryos, but they no longer develop properly, which may be due to the severity of the mutation accumulation they have experienced.

### Transposable element expansion

Barbara McClintock speculated on the genomic stressors associated with plant tissue culture^47^, and transposable element (TE) activation in these conditions is well documented^17,48,49^. Yet until the advent of long-read sequencing, the ability to comprehensively analyze genome-wide TE expansion was limited. Leveraging PacBio HiFi and our haplotype-resolved assembly, we investigated *de novo* TE expansion in the clones.

Analysis of *de novo* structural variants in the long-read samples revealed enrichments of large insertions in the E HiFi somatic embryo around 900 bp and 5500 bp in length, along with a lack of insertions in the trees (Fig. 3a). These high frequency insertions were termed 900 class and 5500 class respectively. The 5500 class was composed of two main sequences, both identified as Copia/LTR transposons, while the 900 class was composed of one sequence with no readily identifiable domains (Fig. 3b, Supplementary Fig. 11 and 12). In both classes, insertions were accompanied by target site duplications of ~10 bp (Supplementary Fig. 13 and 14).

**Fig. 3.**
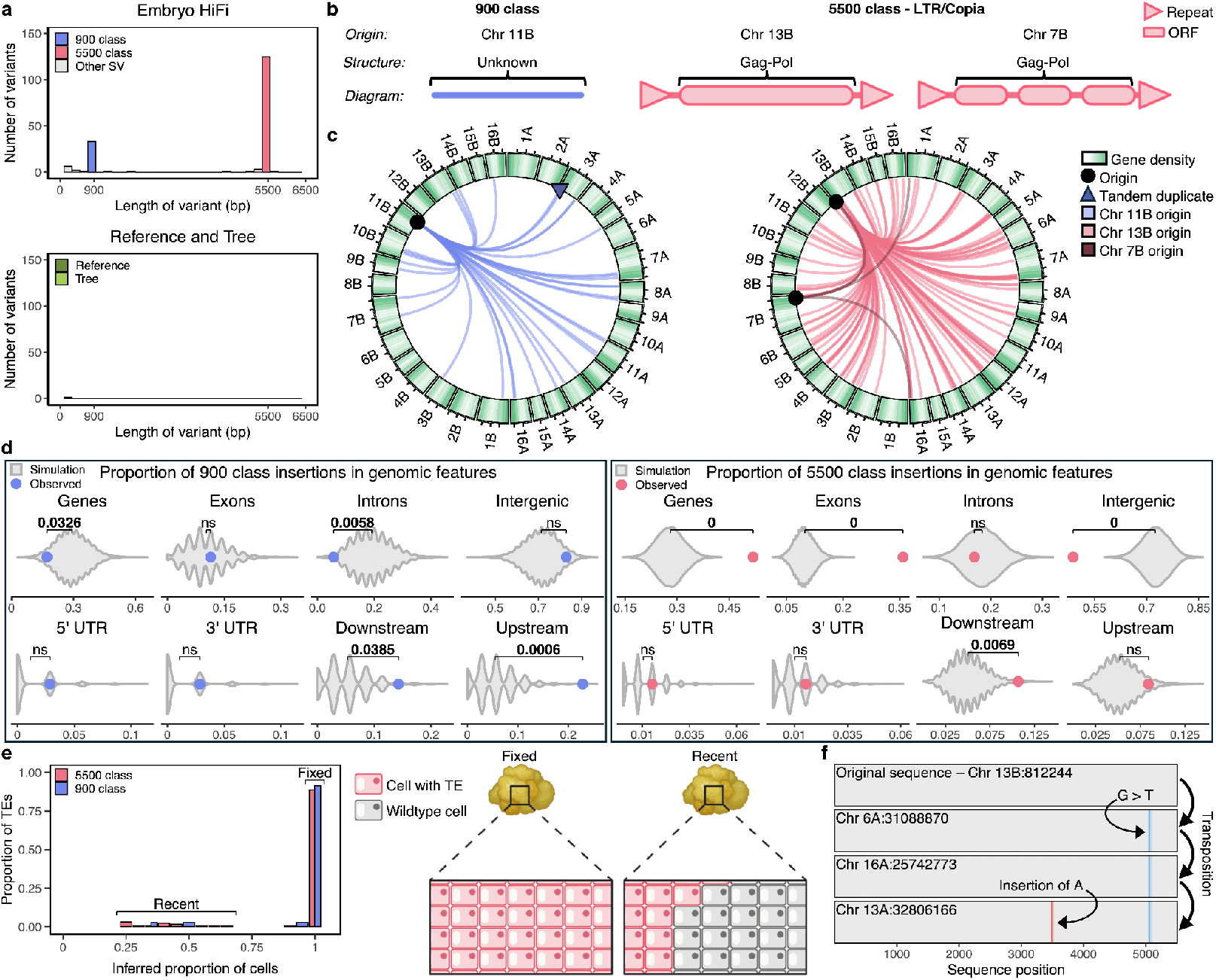
Transposable elements are continuously active in the long read somatic embryo. **a**, Frequencies of *de novo* insertions in the clones sequenced with PacBio HiFi. **b**, Morphology of class 900 and 5500 transposons. **c**, Circos plots depicting transposition of 900 and 5500 class TEs. Lines trace TE movement in the somatic embryo from the putative origin to *de novo* insertion sites. **d**, A comparison of the observed location of TE insertions relative to genomic features. Purple points indicate the observed means of the 900 class TEs and pink points indicate the observed means of the 5500 class TEs. Grey violins represent the distribution of means of 10,000 random simulations. Brackets are labeled with statistically significant differences (p < 0.05) between observed and simulated means. **e**, Class 900 and 5500 TEs observed in different proportions of cells inferred from allele frequency that compose the somatic embryo, accompanied by a diagram. **f**, Tracking a class 5500 TE through multiple transposition events using mutation accumulation.

The putative origins of these insertions were identified based on sequence identity in the haplotype-phased reference assembly, and the transpositions in the somatic embryo were tracked across the genome. There were 35 new 900 class insertions and 131 new 5500 class insertions (Fig. 3c). We found significant bias in insertion sites for both elements, with 900 class insertions enriched in regions up- and downstream of gene bodies and underrepresented within genes and introns (Fig. 3d). The 5500 class insertions showed an enrichment within genes, exons, and up- and downstream of gene bodies, but were underrepresented in intergenic space (Fig. 3d). The insertion bias patterns of the 5500 class are consistent with known patterns of insertion exhibited by Copia/LTR^50^ retrotransposons. We did not find these same biases for non-TE-like insertions (Supplementary Fig. 15a-c). Purifying selection would be expected to remove genic TE insertions, therefore it is unlikely to explain the observed patterns of TE bias. Conversely, SBS and InDels are reduced in genic sequences, consistent with previous findings on somatic mutation in both plants and humans^29,30,36,51^ (Supplementary Fig. 8a).

To assess whether TEs are continuously active in the somatic embryo, TE insertions were called against the haplotype-phased assembly, allowing us to directly infer the proportion of mutated cells. While most TE insertions are present in all cells (100% allele frequency), we detected phased sites with insertions that exist within only a fraction of cells, a result consistent with ongoing TE activity (Fig. 3e, Supplementary Fig. 15d). We also observed sequential transposition through the accumulation of novel mutation, indicating a TE insertion, followed by a mutation event, then subsequent insertion (Fig. 3f, Supplementary Fig. 16 and 17).

Our results show that various transposable elements are continuously active in the somatic embryo. The 900 class insertions exhibit a lack of identifiable motifs, but retain signatures of target site duplication. This suggests they are non-autonomous, though we found no evidence of the corresponding autonomous element and the 5500 class does not share sequence similarity with the 900 class.

Increased transposon activation may reduce the efficacy of using somatic embryos in plant biotechnology, as continual TE mutagenesis may break genes or alter their expression^52^.

### Somatic mutation reveals development

Previous work in plants has leveraged somatic mutation to reveal developmental dynamics^53^, and next generation sequencing has clarified questions about human ontogenesis^20,54^. Using somatic mutation, we explored the developmental underpinnings of clone variability.

The meristems of flowering plants are organized into three distinct cell layers termed L1, L2, and L3^55–57^, with mutations typically restricted to individual layers^21,23^, though the boundary between L2 and L3 is notably less stable than the boundary between L1 and L2^57–63^. Somatic embryogenesis in plants can occur from a single cell^64,65^, which would lead to mutations detected in all cells of the newly generated somatic embryo.

Mutations were called against the phased assembly and restricted to *de novo* SBS and InDels unique to individual clones. As with the transposable element analysis, this allowed us to directly infer the fraction of cells carrying *de novo* mutations. The frequency of these mutations revealed two distinct patterns. Most mutations in the somatic embryos are present in all cells, while some mutations are present at low frequency (Fig. 4a). The high number of unique mutations fixed at 100% allele frequency reveals that single-cell bottlenecks are likely common in the somatic embryos. We hypothesize these bottlenecks occur when the somatic embryos bud (Fig. 4b), and the low frequency mutations that are not present in all cells suggests this process occurs often.

**Fig. 4.**
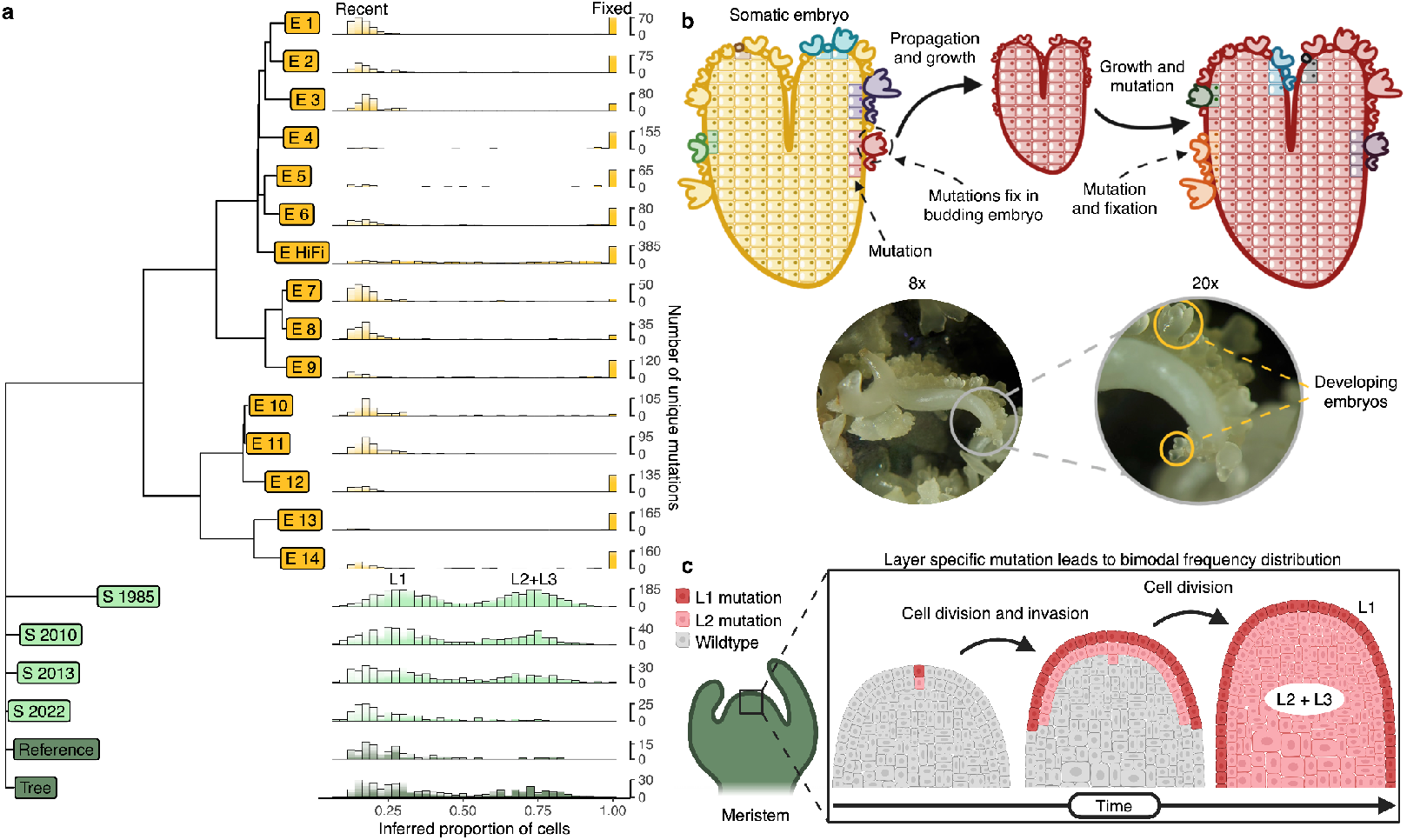
Development and anatomy contribute to differences in mutation burden between clones. **a**, Number of singleton *de novo* SBS and InDels occurring in a proportion of the cells inferred from allele frequency composing each clone. Plots were generated from minimum quality scores of 5-40 and plotted on top of one another with alpha = 0.1. Confidence in the mutation count is therefore depicted by bar darkness, with darker and lighter regions representing higher and lower confidence, respectively. **b**, Diagram and microscopy depicting somatic embryo growth and development. A proposed mechanism of somatic mutation fixation in the somatic embryos is presented in the diagram, and somatic embryos at different developmental stages are labeled in the microscope image, **c**, Schematic portraying a proposed mechanism of mutation fixation in the shoot culture and field-grown tree clones.

Conversely, we found essentially no fixed mutations in the shoot cultures and tree clones, which instead exhibit a distinct bimodal distribution of somatic mutation (Fig. 4a). This pattern would be expected from bulk sequencing of two developmentally distinct sources, like plant cell layers. The impermanence of the boundary between the L2 and L3 layers allows for the propagation of a mutation in both layers, therefore constituting a higher fraction of cells in the organism than mutations constrained to L1 (Fig. 4c).

Somatic mutation in long-term clonal material provides a valuable resource to study anatomy and development in plants. The fixation of mutations in the embryos confirms frequent single-cell bottlenecks, while the low frequency mutations shed light on the ongoing nature of this process. We were also able to rediscover long-studied anatomy of flowering plants by tracking somatic mutations across a haplotype-phased genome assembly. These developmental differences are likely important factors underlying the mutation variation among clonal propagation methods.

## Conclusion

Given the widespread use of cloning in nature, agriculture, and biotechnology, plants represent one of the most critical systems for understanding the dynamics of somatic mutation and its consequences.

Tissue culture is a critical part of genome engineering in many plants, where methodological advancements have largely prioritized increased transformation and editing efficiencies. Our findings suggest another critical area of optimization: minimizing mutation accumulation to preserve the genomic integrity of the plant. Somatic embryos are often used for genome editing because they avoid chimerism, however we find that they exhibit a high mutation rate and extreme genomic instability. The focus on editing efficiency may unintentionally select for systems with higher mutation accumulation.

These somatic embryos have been maintained in culture for several decades, and there is evidence of continual mutation accumulation over time. We predict that the extreme mutation burden we observed could be ameliorated in material subject to shorter *in vitro* periods, however we also find potential evidence of whole-chromosome duplication occurring during the initial stages of culture. Sequencing regenerated material to facilitate the early detection of potentially deleterious mutations may be advised to avoid advancing and propagating sub-optimal clones. Future research should identify mutation over shorter timescales after initial embryogenesis and under different culturing conditions. While somatic mutations may impact the viability of the plants, they are exceedingly unlikely to have any effect if consumed.

In addition to these practical implications, our observations address fundamental questions about mutation in plants. We identified distinct mutational differences among clonal material, from single base changes to whole-chromosome duplications. These discoveries inspire future work to investigate how differences in DNA damage, repair, and development drive somatic mutation dynamics through time and space in plants.

## Methods

### Somatic embryo initiation & maintenance

A repetitively embryogenic somatic embryo culture derived from anther tapetal tissue of *Juglans regia* cultivar ‘Chandler’^26^ was maintained on corrected Driver and Kuniyuki Walnut (DKW-C) medium^66^ without growth regulators, supplemented with 30g/L sucrose, and solidified with gellan gum. Cultures were kept on Petri plates continuously in the dark at room temperature. As new embryos developed by direct embryogenesis^67^, a subset of embryos was transferred to fresh medium approximately weekly since 1995.

### Shoot culture introduction & maintenance

Microshoot cultures of ‘Chandler’ were established by nodal cuttings taken from the most original field-grown source in 1985, 2010, 2013, and 2022^68^. The 2010, 2013, and 2022 cuttings were generated from the same tree that provided material for the new reference assembly. Cultures were maintained under 16 hour light on DKW-C medium supplemented with 1 mg/L BAP and .01 mg/L K-IBA^66^ with transfer to fresh medium every 2-4 weeks^69,70^. Recent work in strawberry suggests increased hormone levels associated with micropropagation led to higher rates of mutation^71^.

### DNA and RNA extraction

One somatic embryo from each subpopulation (12 in total) was selected at random and divided in half. DNA was extracted from one of the halves using the DNeasy Plant Mini Kit (QIAGEN) and RNA was extracted from the remaining half using the RNeasy Plant Mini Kit (QIAGEN). DNA was also extracted from two randomly selected embryos from the same subpopulation with the same kit. Two embryos from the same subpopulation located next to one another were pooled and nuclei were extracted using the Circulomics Plant Nuclei Protocol. Pooling was done to meet minimum tissue requirements for extraction and reduce variability, as the adjacent embryos from the same subpopulation are likely derived from the same progenitor. High molecular weight DNA was extracted from the nuclei using the Nanobind HMW DNA Extraction - Plant Nuclei kit from Circulomics. Shoot apical meristem, stem, leaf, and petiole tissue were collected from each of the shoot cultures and DNA was extracted using the DNeasy Plant Mini Kit (QIAGEN). Leaf tissue was collected from across the canopy of the two field-grown trees. Nuclei were isolated from both samples using the Circulomics Plant Nuclei Protocol, and high molecular weight DNA was extracted using the Nanobind HMW DNA Extraction - Plant Nuclei kit from Circulomics.

### Library prep and sequencing

All library preparation and sequencing was conducted by the UC Davis Genome Center DNA Technologies and Expression Analysis Core Facility. Illumina Whole Genome Sequencing (150 bp, paired-end) was performed using the Illumina NovaSeq S4 300 platform. Poly-A RNA sequencing (150 bp, paired-end) was conducted using the Illumina NovaSeq X 25B platform. High molecular weight DNA was sequenced on the PacBio Sequel II platform (Supplementary Table 1).

### Omni-C data generation

Leaf tissue from across the canopy of the Reference clone was collected, pooled into a 50 ml tube, and flash frozen with liquid nitrogen. The sample was shipped to Cantata Bio for Omni-C library preparation and sequencing. The Omni-C library was prepared using the Dovetail® Omni-C® Kit according to the manufacturer’s protocol. Briefly, the chromatin was fixed with disuccinimidyl glutarate (DSG) and formaldehyde in the nucleus. The cross-linked chromatin was then digested in situ with DNase I. Following digestion, the cells were lysed with SDS to extract the chromatin fragments and the chromatin fragments were bound to Chromatin Capture Beads. Next, the chromatin ends were repaired and ligated to a biotinylated bridge adapter followed by proximity ligation of adapter-containing ends. After proximity ligation, the crosslinks were reversed, the associated proteins were degraded, and the DNA was purified then converted into a sequencing library using Illumina-compatible adaptors. Biotin-containing fragments were isolated using streptavidin beads prior to PCR amplification. The library was sequenced on an Illumina HiSeq X platform to generate 400 million 2 × 150 bp read pairs.

### Reference genome assembly

To allow better mutation detection^22,72^, a haplotype-phased reference genome assembly was constructed using Hifiasm v0.19.8^73–75^ from the HiFi CCS reads and Omni-C reads generated for the field grown Reference clone. The primary and phased contigs generated by Hifiasm were then scaffolded using the previous Oxford Nanopore Technologies, Hi-C, and short read generated walnut cultivar ‘Chandler’ genome (‘Chandler’ v2.0) with RagTag v2.1.0^76^.

A ‘Chandler’ genetic map was constructed using genotyping-by-sequencing data from 436 self-pollinated ‘Chandler’ progeny. Data were imputed using FSFHap^77^, resulting in a dataset of 16,282 SNPs. One individual with >5% missing data was discarded. SNPs were filtered to retain only those with 30-70% heterozygous genotypes and <5% missing data, resulting in the removal of 37 SNPs. Testing for segregation distortion of genotypes and alleles revealed no remaining SNPs with chi-square p-values below 0.001. The ASMap package in R^78^ was used to construct a genetic map of 16,245 markers and 3,660 bins and near-perfect agreement in marker order between genetic and physical maps (Supplementary Fig. 6). The genetic map was then used to verify phasing in the reference genome by assessing the accuracy of the haplotype assignment by Hifiasm by identifying haplotype and primary assembly correspondence with SyRI v1.5.4^79^ and using custom R scripts (Supplementary Fig. 1b).

The assembled haplotypes were concatenated into a single fasta file, creating four assemblies: The primary assembly, haplotype A assembly, haplotype B assembly, and a concatenated haplotype A and B assembly.

Omni-C reads were then aligned to the primary, haplotype A, and haplotype B assemblies and contact matrices were generated using the Dovetail Genomics protocol (https://omni-c.readthedocs.io/en/latest/contact_map.html) and JuicerTools^80^ (Supplementary Fig. 2). Telomere presence was assessed using the R package ggenomics (https://github.com/matthewwdavis/ggenomics).

### Annotating the genome assemblies

For the primary, haplotype A, and haplotype B genome assemblies, annotations were generated using Liftoff v1.6.3^81^ and the ‘Chandler’ v2.0 GFF^28^.

Transposable elements (TEs) were annotated on the primary assembly using The Extensive de novo TE Annotator (EDTA) v2.1.0^82^ with default parameters. We removed transposable elements that overlapped with the filtered plant protein database using protExcluder v1.2 (https://www.canr.msu.edu/hrt/uploads/535/78637/ProtExcluder1.2.tar.gz) to reduce the exclusion of genes in any further gene prediction. The TEs were re-annotated on the primary contig assembly using a filtered, non-redundant TE library, and the assembly was softmasked.

The annotations were further processed for downstream analysis in R v.4.4.2^83^ with custom scripts. Annotated genes and CDS were extracted, transcribed, and translated. Genes and CDS that did not begin with a start codon (AUG) and end with stop codons (UAA, UAG, UGA) were removed. Intergenic space, 5’ UTRs, 3’ UTRs, and introns were annotated with custom R scripts.

### Assessing genome assemblies

To assess assembly completeness, BUSCO v5.7.1^84^ analysis was performed using the eudicots_odb10 database. Reference tree reads were assessed for 21-mer content using jellyfish v2.2.10^85^ and the GenomeScope v2.0^86^ web browser tool was used to assess the kmer distribution. The annotated genes in the GFF were used to determine the proportion of the genome constituting gene bodies (Supplementary Fig. 3). Assembly statistics were evaluated using SeqKit v2.10.0^87^ (Supplementary Fig. 1, Supplementary Table 2).

### Comparing Assemblies

Chromosome–scale assemblies were aligned using Minimap2 v2.17-r941^88,89^ and structural variation between the assemblies was assessed using SyRI and visualized using plotSR v0.5.4^90^. Structural similarity was also assessed between the assemblies using the Minimap2 generated alignments and dot plots generated by the D-GENIES^91^ browser tool (Supplementary Fig. 4).

### Genome mappability

Mappability was assessed using GenMap v1.3.0^92^, specifying a kmer length of 150 and allowing for 0 errors. When annotating regions as mappable using custom R scripts, we retained regions with a mapping probability of 1.

### Calling SBS and small InDels

Short read quality was assessed using FastQC v0.11.9^93^ and reads were trimmed using Trimmomatic v0.39^94^. Trimmed reads were then aligned to the primary reference assembly and the concatenated haplotype assembly using Minimap2 v2.17-r941. The resulting BAM files were sorted, duplicates were marked, and the files were indexed with SAMtools v1.13^95^. SBS and InDels were called against the primary and concatenated genome assemblies using DeepVariant v1.6.1^96^ with the model type ‘WGS’ to generate VCF and GVCF files. For regions within the GVCF that contained no variants, median depth of the region was reported.

Long reads were aligned to the primary assembly and the concatenated haplotype reference assembly using the PacBio Minimap2 wrapper pbmm2 v1.13.1 (https://github.com/PacificBiosciences/pbmm2) with the preset ‘HIFI’. The resulting BAM files were sorted and indexed using SAMtools v1.13. SBS and InDels were called against the primary and concatenated assemblies using DeepVariant v1.6.1 with the model type ‘PACBIO’ to generate VCF and GVCF files. For regions within the GVCF that contained no variants, median depth of the region was reported.

The VCFs were additionally annotated in R with custom scripts. Sites were assessed for mapping probability. Reference allele and alternate allele depths were extracted, and the total depth of a site was recalculated by adding the reference allele depth and the alternate allele depth. The variant allele frequency of a site was recalculated by dividing the alternate allele depth by the recalculated total depth. Median depth was calculated across individuals at the same position, as was median quality.

### Verifying samples are clonal

SBS and small InDels were called against the ‘Chandler’ primary assembly using WGS data of 5 other *Juglans regia* cultivars (Franquette, Hartley, Payne, PI159568, & Waterloo). A SBS PCA was generated using R to visualize the relationship between the clones and the other cultivars, as well as the relationship of the clones to each other. (Supplementary Fig. 18).

### Calling structural variants

The sorted and indexed BAM files generated from aligning long reads to the concatenated haplotype assembly with pbmm2 were used to discover and call structural variants (SVs) using pbsv v2.9.0 (https://github.com/PacificBiosciences/pbsv) and generate structural variant VCFs (svVCFs).

### Mapping RNA sequencing data

RNA sequencing data was pseudo-aligned to the primary reference assembly and counts were generated and normalized to transcripts per million (TPM) using kallisto 0.51.0^97^.

### Identifying *de novo* mutations

To identify small-scale *de novo* mutation and ancestral heterozygosity, all primary assembly sample VCFs were organized into groups based on tissue type and propagation method. All somatic embryos were classified as “embryo” group, all shoot cultures were classified as “shoot” group, and all field-grown trees were classified as “tree” group. Variants that were found in more than one group were considered to be ancestral heterozygosity. Variants found only within the “embryo” group were determined to be *de novo* mutations, as all embryos are derived from one initial embryo, and removing mutations shared by the embryos would be removing real *de novo* mutations. Variants found between members of the same group in the “shoot” and “tree” groups were removed, as these propagations were independent of one another and shared mutation was likely ancestral heterozygosity. The remaining mutations in these individuals were determined *de novo* (Supplementary Fig. 7). A similar assessment of the combined haplotype VCFs was performed. In this case, any variants shared by more than one group is dubious, as heterozygosity should be accounted for in the assembly since both haplotypes are represented. In this case, any shared variants between the groups are removed. SnpEff v5.1d^98^ was used to evaluate the effect of mutations.

To classify *de novo* structural variation, the three svVCFs were assessed for shared variants. If a variant was present in more than one sample, it was determined ancestral. If it was only present in a single sample, it is considered *de novo*.

### Generating somatic mutation phylogeny

Phylogenies were constructed from *de novo* somatic SBS and InDels using the R packages ape^99^, phangorn^100^, ggtree^101^, and polymorphology2 (https://github.com/greymonroe/polymorphology2). Mutations were filtered for sites in regions with a mappability score of 1, a median quality score >=30, and a median depth >=15. Distance between individuals was calculated using the JC69 model^102^ and clustering was done using the unweighted pair group method with arithmetic mean (UPGMA^103^). The phylogenies were bootstrapped 10,000 times and branch lengths were made proportional to the number of unique mutations in a branch.

### *De novo* SBS tricontext

Previously identified *de novo* SBS were filtered for a median site quality >=30, median depth >=15, and a mappability score of 1. The number of SBS occurring in each context was counted, then corrected by the number of times the trimer occurred in the reference.

### Identifying mutation location

Mutations defined as *de novo* with a median site quality >=30, median depth >=15, and a mappability score of 1 were assessed for overlap with the curated list of genes using custom R scripts and the polymorphology2 package.

### Calculating scaled depth in chromosomes

Genomic instability can have dramatic effects on phenotype^104,105^. All sites previously identified as ancestral heterozygosity when aligned to the primary reference genome were assigned haplotypes using the genetic map. The alternate call was used to represent one haplotype, with the allele frequency and alternate read depth associated with it. The other haplotype was then represented by the reference call, with one minus the allele frequency representing the frequency at that site and the reference depth representing the read depth. The depth for each site was scaled by the mean depth of the sample. Each chromosome was separated into 7 equally sized segments, and the mean of the scaled depth was taken at each segment. Haplotype assigned scaled read depth was also used without windows to plot chromosome duplications, somatic recombinations, and large deletions (Supplementary Fig. 9a).

### Counting telomeric repeats

To identify telomeric repeats in the samples, the reads aligning to the nuclear chromosomes from the bam files previously generated were extracted for each sample using SAMtools v1.13. 21-mers were counted in the nuclear reads using jellyfish v2.2.10. The counts for the telomeric repeat (TTTAGGG) occurring 3 times in a row were extracted and divided by the total number of base pairs in the nuclear reads for the sample.

### Comparing telomeric repeat numbers

The telomeric repeat counts of all embryos were compared to the counts of shoots and trees with a Welch’s t-test using custom R scripts. Embryos were then separated into the two major clades and compared to one another with a Welch’s t-test. The clade with fewer repeats was then compared to the trees and shoots in a similar manner.

### Assessing allele frequency in aneuploids

The ancestral sites that were previously phased were used for assessing allele frequency. Each chromosome was separated into 250 equally sized windows, and the mean allele frequency of each window was calculated. The sum of all haplotype A and haplotype B depths in each window was also calculated. A chi-square test was performed to determine if the sum of haplotype A and B depths differed significantly from the expectation of 0.5 and 0.5, the expected proportion of depths at a heterozygous site in a diploid.

### Simulating ploidy changes

Each site in the VCF of the clones sequenced with PacBio HiFi was filtered for a quality >=40 and a depth >=30. The difference in sequencing depth between the depth of the reference allele and alternate allele of each site was then simulated with a weighted binomial distribution. To simulate a diploid, the weighted depths were ½ and ½, a triploid was weighted ⅓ and ⅔, and an asymmetric tetraploid was weighted ¼ and ¾. These distributions for each chromosome were plotted, with the observed distributions of the chromosomes also plotted (Supplementary Fig. 10b).

### Timing duplications

The relative timing of the chromosome 4A and 9B duplications was identified through allele frequencies. In the primary and haplotype-resolved assemblies, *de novo* mutations were filtered for those shared by all embryos and with a median quality >=30, depth >=15, and a mapping probability of 1. A chi-square test was performed to determine if the mutations differed significantly from the diploid expectation of 0.5 and 0.5. If the p-value was <0.01 and the allele frequency was >0.5, the site was determined as duplicated. If allele frequency was <0.5, it was non-duplicated. The mutations occurring on chromosomes 4 and 9 were plotted as a scatterplot to visualize. The duplicated mutations on these chromosomes were corrected for chromosome length and compared with Welch’s t-test. The ratios of the duplicated to non-duplicated mutations on these chromosomes were also compared using Welch’s t-test (Supplementary Fig. 9c-e).

### Determining chromosome loss

To identify if the chromosome 4 duplication occurred multiple times independently or occurred once and was lost, *de novo* mutations in the haplotype-resolved assemblies in chromosome 4 were filtered for quality >=30. A chi-square test was performed to determine if the mutations differed significantly from the expectation of 0.5 and 0.5 and a False Discovery Rate (FDR) correction was applied. If the p-value was <0.05 and the allele frequency was >0.5, it was determined to be a mutation that occurred before duplication. If the allele frequency was <0.5 it was identified as a post-duplication mutation. The mutations that occurred post-duplication, but were still found in the non-duplicated samples were identified as retained post-duplication mutations (Supplementary Fig. 9b).

### Comparing expression across chromosomes

RNA count data were filtered for TPM >1.0 and the median TPM was calculated across each chromosome. The mean of the median chromosome TPMs was then calculated for each sample, and the median chromosome TPM was normalized by the sample mean TPM.

### Gene-level differential expression

Somatic embryo samples with RNA evidence were investigated for differential RNA expression. The TPM of each transcript in every clade was compared to the TPM of that transcript in embryos that were not within the clade using a Welch’s t-test. Resulting p-values were corrected using the FDR correction and adjusted p-values <0.05 were considered significant. All adjusted p-values were −log10 transformed.

### *De novo* structural variation

To identify *de novo* structural variation, the three samples with svVCFs generated (Embryo HiFi, Reference, Tree) were filtered for SVs that only existed in a single sample.

### Identifying transposition and origin

Using Nucleotide BLAST v2.15.0+^106^, all *de novo* SVs detected were used as query sequences against the concatenated haplotype-phased reference assembly. Sequences were then assessed for origin location found in the haplotype-phased assembly. The structural variant sequences were grouped based on insertion length and origin location into two main categories, 900 class and 5500 class variants. The first and last half of a 1792 bp insertion matched the 900 bp class origin location, indicating an insertion followed by a tandem duplication. The sequences within classes were aligned to one another using MUSCLE^107^ within the R package msa^108^. The alignments were visually inspected with Geneious Prime 2022.2.1 (https://www.geneious.com) and manually trimmed to remove the target site duplication sequence incorporated during transposition^109^. Using Geneious Prime, the necessary sequences were reverse complemented and all sequences were realigned using MAFFT^110^ (Supplementary Fig. 13 and 14).

### Aligning identified TEs for consensus

Verified and trimmed TE sequences were aligned to one another using MUSCLE in the R package msa. The consensus sequence was determined as the most common nucleotide at a specific position. TE sequences with large size differences were removed from the alignment to improve legibility (Supplementary Fig. 16 and 17).

### Assessing transposon morphology

Using Nucleotide BLAST v2.15.0+, all *de novo* SVs detected were used as query sequences against the fasta file of EDTA predicted transposable elements. Open reading frames (ORFs) in the 900 bp class and 5500 bp class of sequences were identified using NCBI ORFfinder (https://www.ncbi.nlm.nih.gov/orffinder/).

Identified ORFs were searched for similarity in the NCBI database using Protein BLAST^106^. Geneious Prime was used to further validate ORFs and identify repeat regions in the 900 bp and 5500 bp classes (Supplementary Fig. 11 and 12).

### Comparing observed insertions to random

Each insertion within the 900 bp and 5500 bp classes was assessed for its location relative to features annotated within the GFF. The insertions were annotated for being within gene bodies, 5’ untranslated regions (UTRs), 3’ UTRs, exons, introns, intergenic space, upstream regions, and downstream regions. Upstream regions were defined as sequence 1000 bp before the annotated gene start and downstream regions were defined as 1000 bp after annotated gene stop. Each insertion was then simulated to land at a random position in the target chromosome 10,000 times and the simulated insertions were assessed for their location relative to annotated features. The mean of the simulated distribution was compared to the mean of the observed data, to calculate significance.

### Proportion of cells containing transposons

Windows were created from allele frequency of 0 to 1 in increments of 0.05. The proportion of 900 bp class and 5500 bp class insertions was assessed in each window.

### Fixation of SBS and InDels

SBS and InDels called from the parsed haplotype-phased VCFs were further annotated for variants that were unique to a single sample. The VCFs were filtered for sites with quality values >=5 and depth >=10. For each sample, histograms were generated to display the occurrences of variant allele frequencies at quality scores 5-40 in increments of 1. The histograms of different quality thresholds were plotted on top of one another at an alpha value of 0.1.

### Diagrams

Diagrams were created using Microsoft PowerPoint v16.95 and BioRender.

## Data availability

All raw sequencing data generated for this project are publicly available through NCBI BioProject PRJNA 1271128. The primary genome assembly and annotation of *Juglans regia* cv. ‘Chandler’ is available through NCBI BioProject 1289367, the haplotype A (hap1) assembly and annotation is available through NCBI BioProject 1289368, and the haplotype B (hap2) assembly and annotation is available through NCBI BioProject 1289370. Assemblies, annotations, VCF files, EDTA output, and verified annotations used for analysis are also available at Figshare in the project Walnut clones (https://figshare.com/account/home#/projects/252110). Code for this research is maintained at https://github.com/matthewwdavis/Walnut-Mutation-Accumulation.

## Acknowledgments

We thank the Monroe and Brown labs, Julin Maloof, Luca Comai, Daniel Runcie, Jeff Ross-Ibarra, John Davis, Detlef Weigel and Kylie Roland for insights and feedback. We also thank Noah Feinberg, and Sriema Lalani Walawage for laboratory advice and assistance. This work was supported by the FFAR Fellows Program. Research was conducted at the University of California Davis, which is located on land that was the home of the Patwin people for thousands of years.

## Author contributions

M.W.D, P.J.B., J.G.M. conceptualized the study, determined appropriate methodology, performed formal analysis, validated results, curated data, and visualized data. M.W.D. wrote the original draft. M.W.D., P.J.B. and J.G.M. reviewed and edited the manuscript. P.J.B. and J.G.M. contributed resources, acquired funding, and supervised the project. C.A.L. maintained clonal cultures. C.L. conducted formal analysis and performed data visualization. E.L. contributed to formal analysis and visualization. M.W.D., L.M., and M.L. contributed to data collection and generation. F.L. contributed to visualization.

## Supplementary Materials

### Supplementary Figures

**Supplementary Fig. 1.**
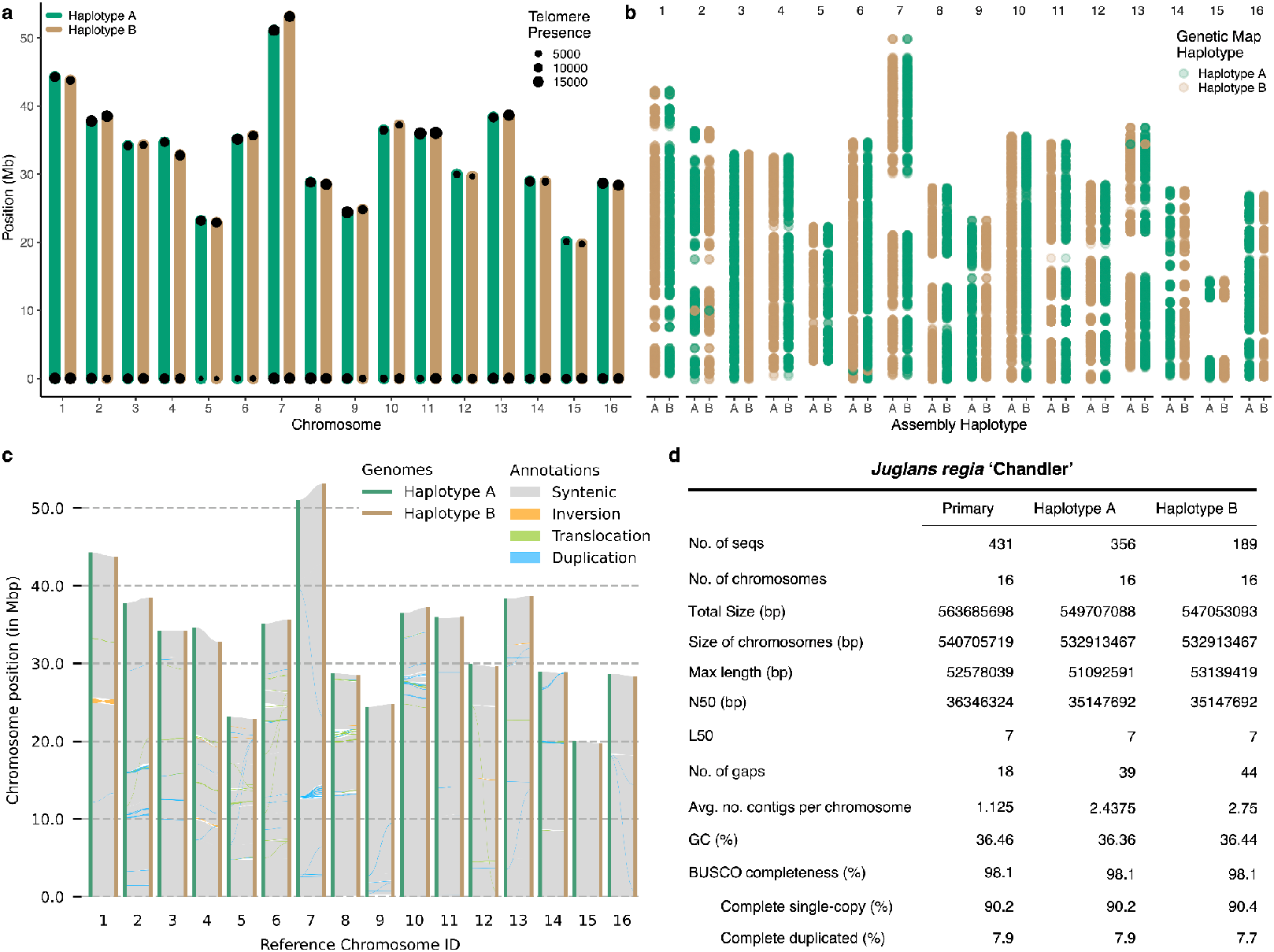
A haplotype-phased *Juglans regia* ‘Chandler’ genome assembly. **a**, Depiction of haplotype phased genome assembly. Bars represent length of chromosome, point size represents location and number of telomeric repeats. **b**, The accuracy of haplotype phasing as determined by the genetic map. The haplotype assigned to the assembly is represented on the x axis and the color of the point corresponds to the genetic map haplotype. **c**, Structural variation between the two haplotypes of *Juglans regia* ‘Chandler’. **d**, Table depicting assembly statistics.

**Supplementary Fig. 2.**
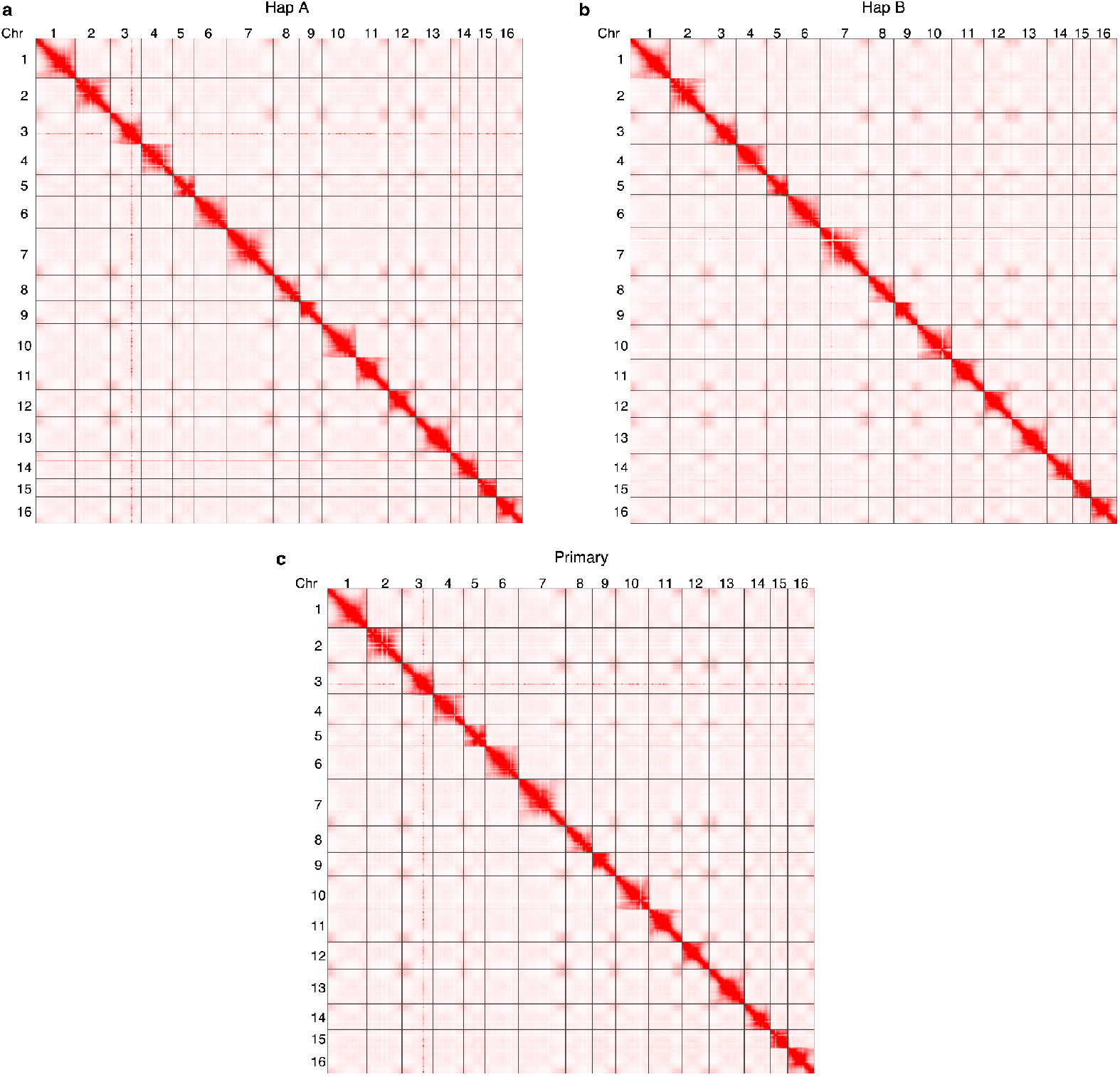
Omni-C contact maps of the three genome assemblies. **a**, Contact map of Omni-C reads to the haplotype A Reference tree genome assembly. **b**, Contact map of Omni-C reads to the haplotype B Reference tree genome assembly. **c**, Contact map of Omni-C reads to the primary Reference tree genome assembly.

**Supplementary Fig. 3.**
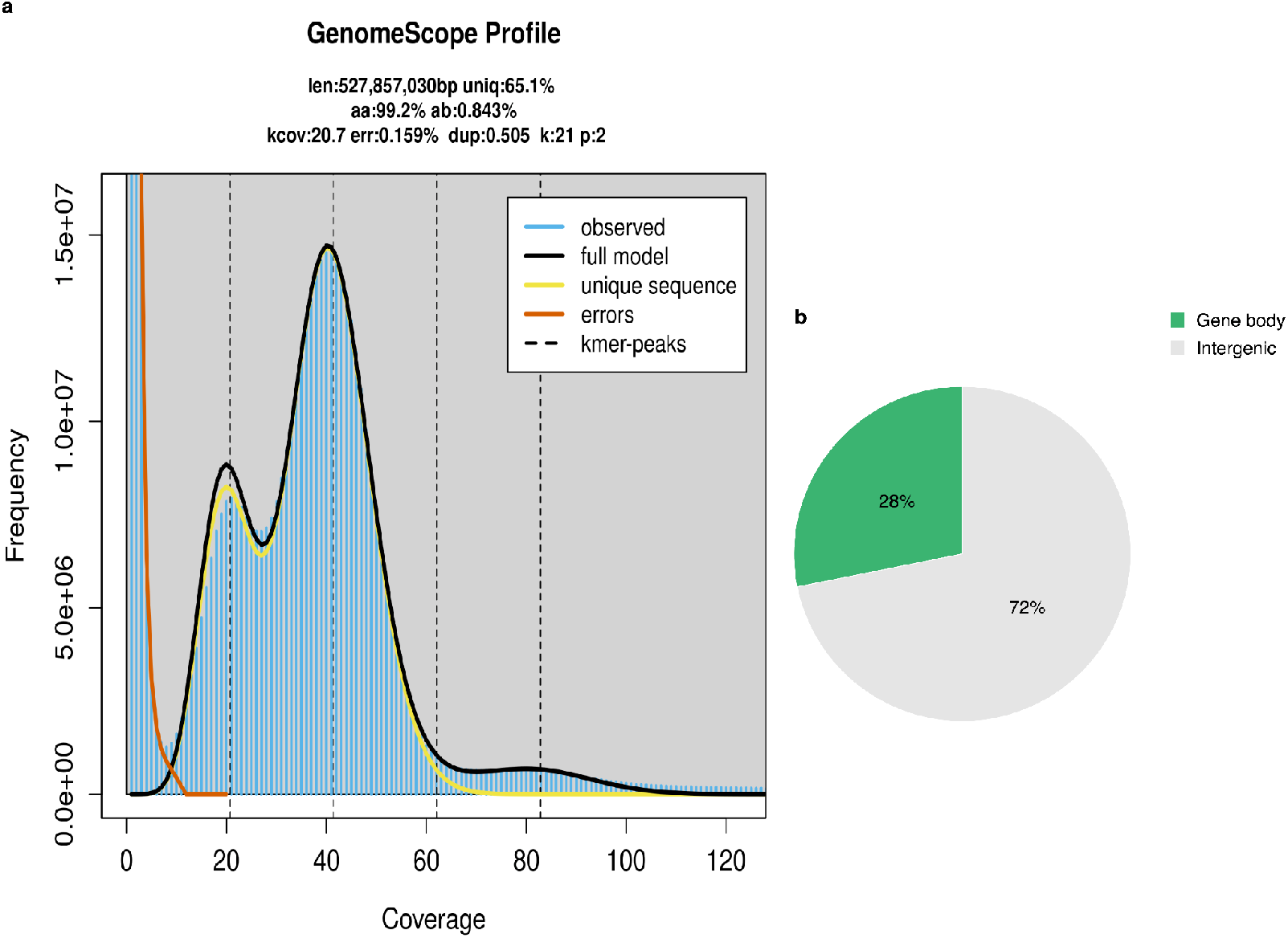
Genomic composition of *Juglans regia* ‘Chandler’. **a**, GenomeScope profile of the reference reads. **b**, Proportion of the genome that is annotated gene bodies compared to the rest of the genome.

**Supplementary Fig. 4.**
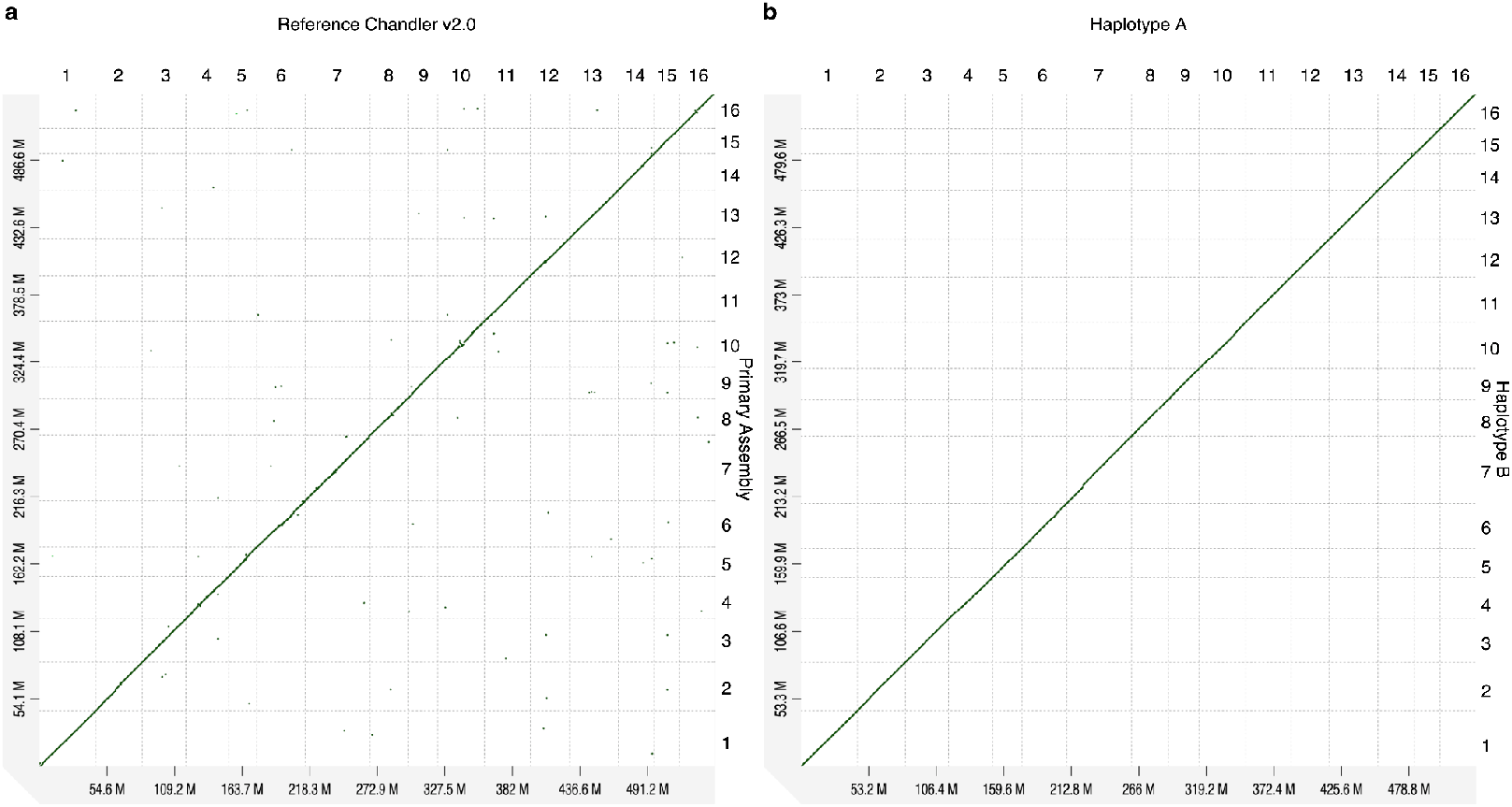
Genome wide dot plots. **a**, D-Genies dot plot visualizing the alignment of the Reference Chandler V2.0 assembly and the newly constructed primary sequence. **b**, D-Genies dot plot visualizing the alignment of the haplotype A assembly and the haplotype B assembly.

**Supplementary Fig. 5.**
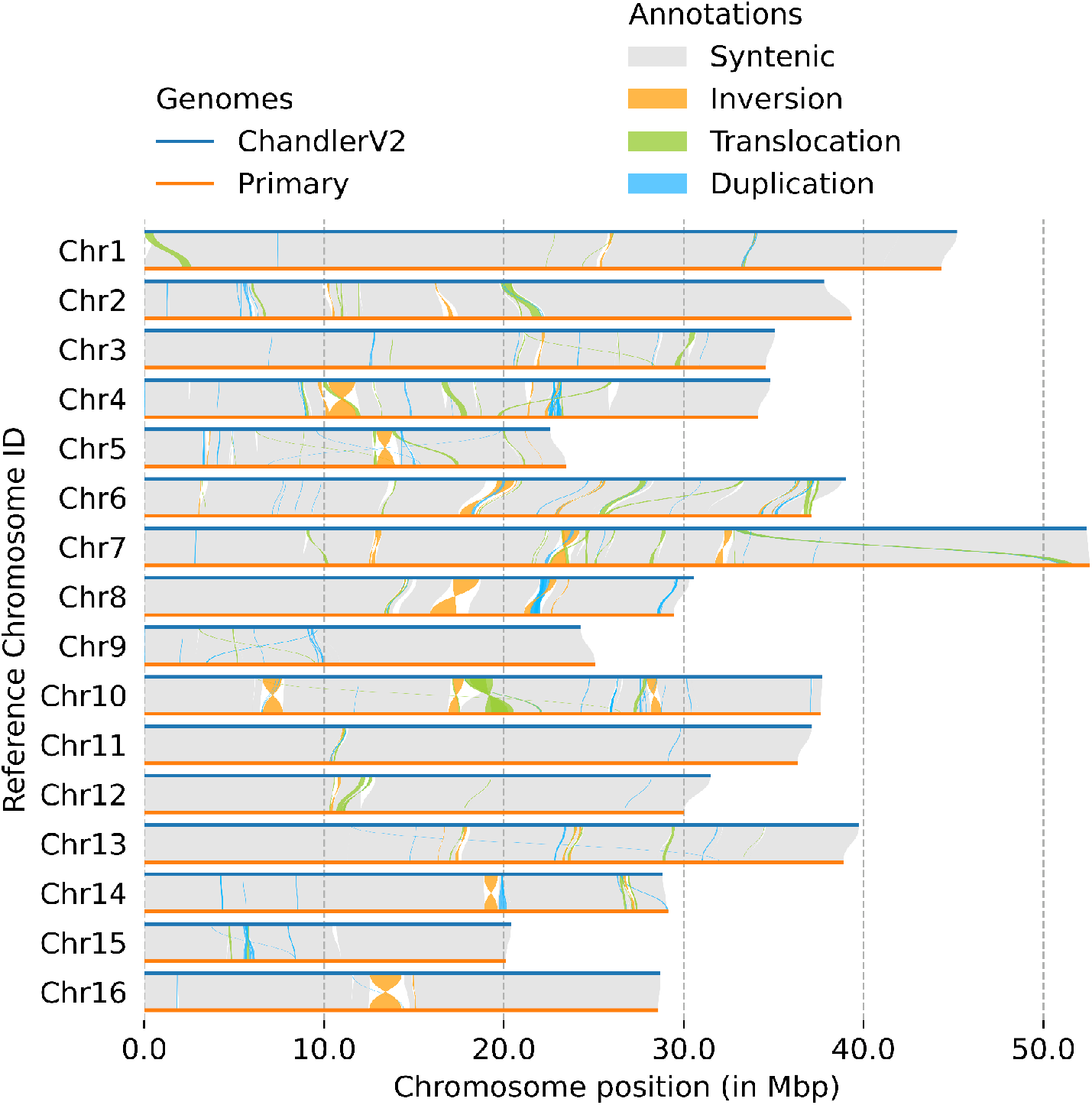
Structural variation between Chandler v2.0 and the primary sequence. Structural variation between the Chandler v2.0 reference sequence and the primary assembly.

**Supplementary Fig. 6.**
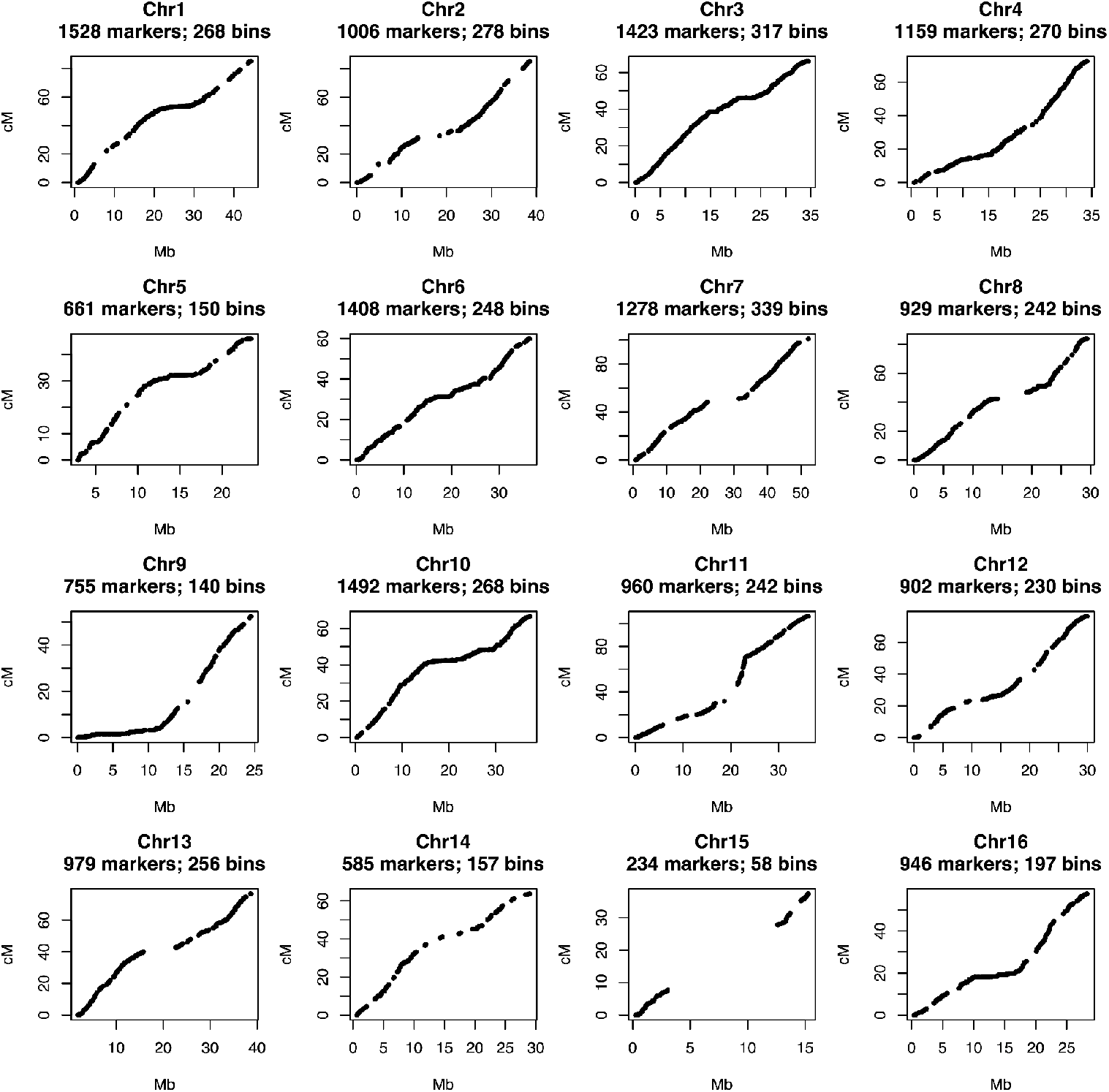
Genetic map of *Juglans regia* ‘Chandler’. The agreement of marker order in the genetic map and physical map for the 16 chromosomes of *Juglans regia* Chandler’.

**Supplementary Fig. 7.**
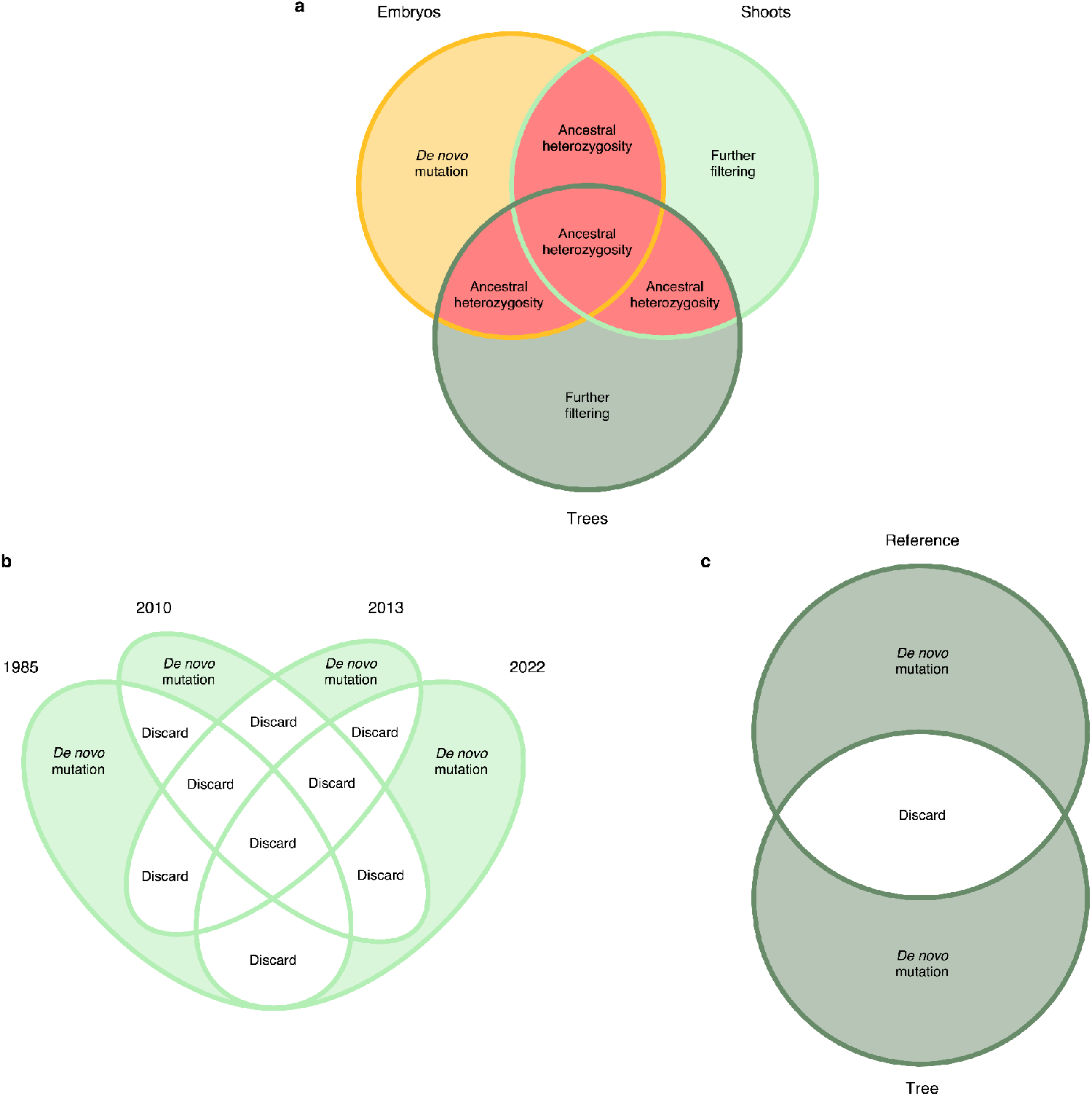
Determining *de novo* mutation and ancestral heterozygosity. **a**, Venn diagram depicting the determination of *de novo* mutation in the somatic embryos and ancestral heterozygosity in the clones. **b**, Venn diagram depicting the determination of *de novo* mutation in the shoot cultures. **c**, Venn diagram depicting the determination of *de novo* mutation in the field-grown trees.

**Supplementary Fig. 8.**
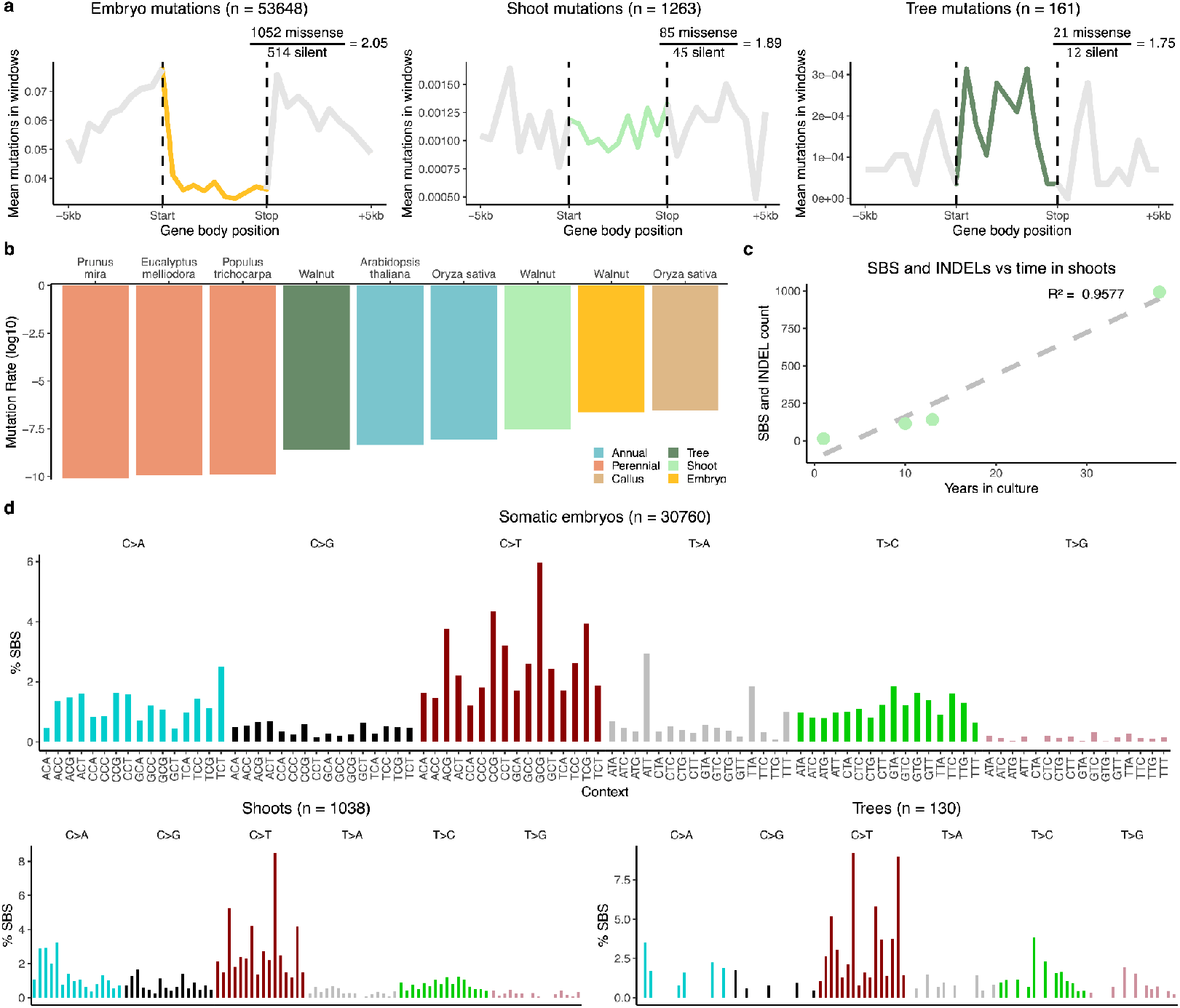
Somatic embryos exhibit a distinct mutational profile. **a**, *De novo* SBS and InDels in the gene body compared to upstream and dovznstream regions. Samples pooled by method of clonal propagation, n value refers to the number of total mutations. **b**, Comparison of previously reported somatic and germline yearly mutation rates in other species and the yearly somatic mutation rates of the walnut clones. **c**, SBS and InDels in shoot clones introduced to tissue culture al different times. The relationship between years in culture and mutation count was assessed with a linear model, and model fit was determined. **d**, *De novo* SBS spectrum of samples pooled by method of clonal propagation. The n value refers to the number of total mutations.

**Supplementary Fig. 9.**
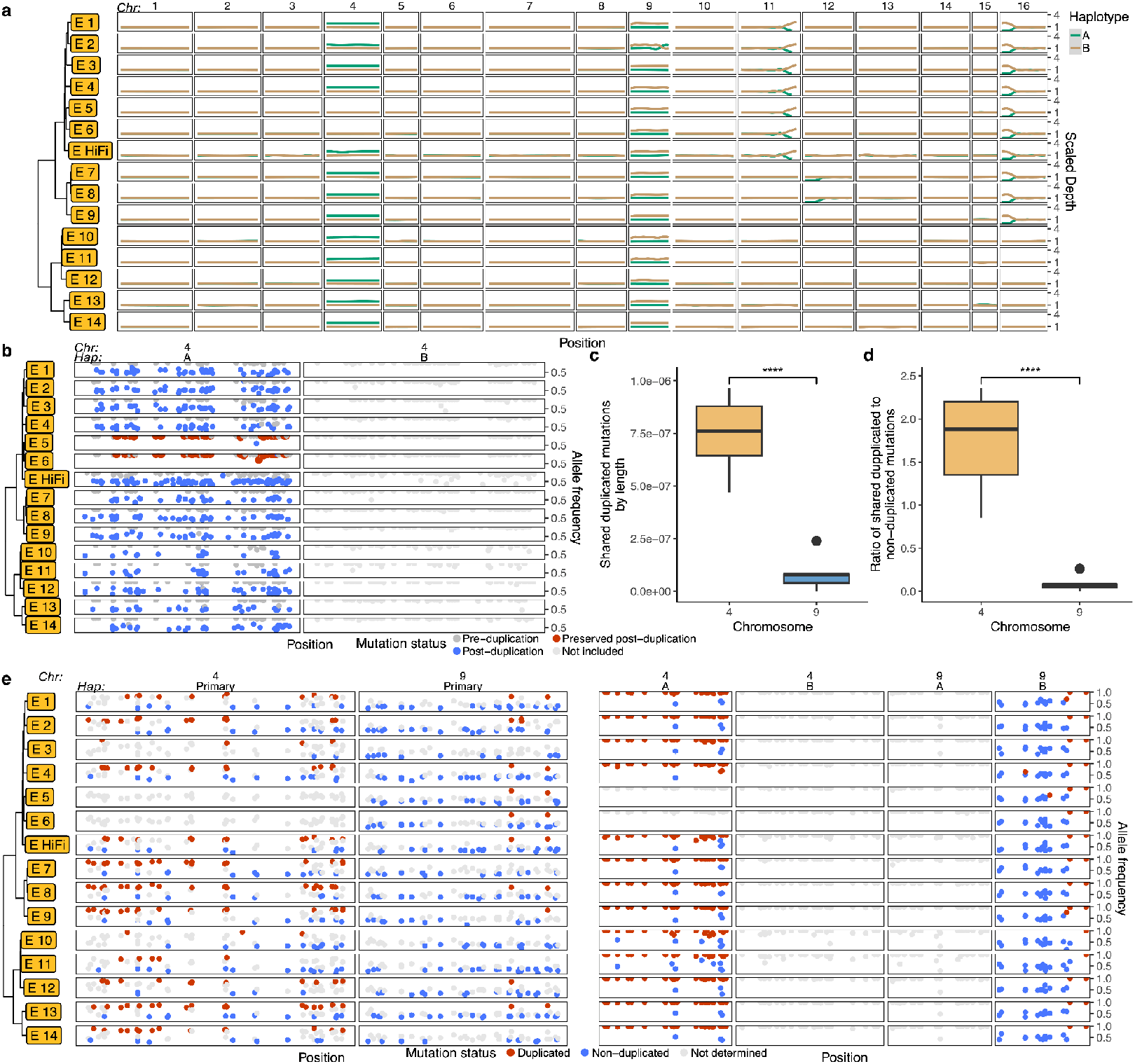
Dating the chromosomal duplications. **a**, Haplotype-resolved depth across chromosomes in the somatic embryos. **b**, *De novo* mutations that occurred after chromosomal duplication (blue) that also exist in the non-duplicated samples (red). **c**, The number of duplicated shared *de novo* mutations corrected by chromosome length in chromosomes 4 and 9. **d**, The ratio of duplicated shared *de novo* mutation to non-duplicated shared de novo mutations in chromosomes 4 and 9. **e**, Shared de novo mutation by all somatic embryos in the primary and haplotype-resolved assemblies. Red points denote mutations that occurred before duplication, blue points are mutations that occurred after.

**Supplementary Fig. 10.**
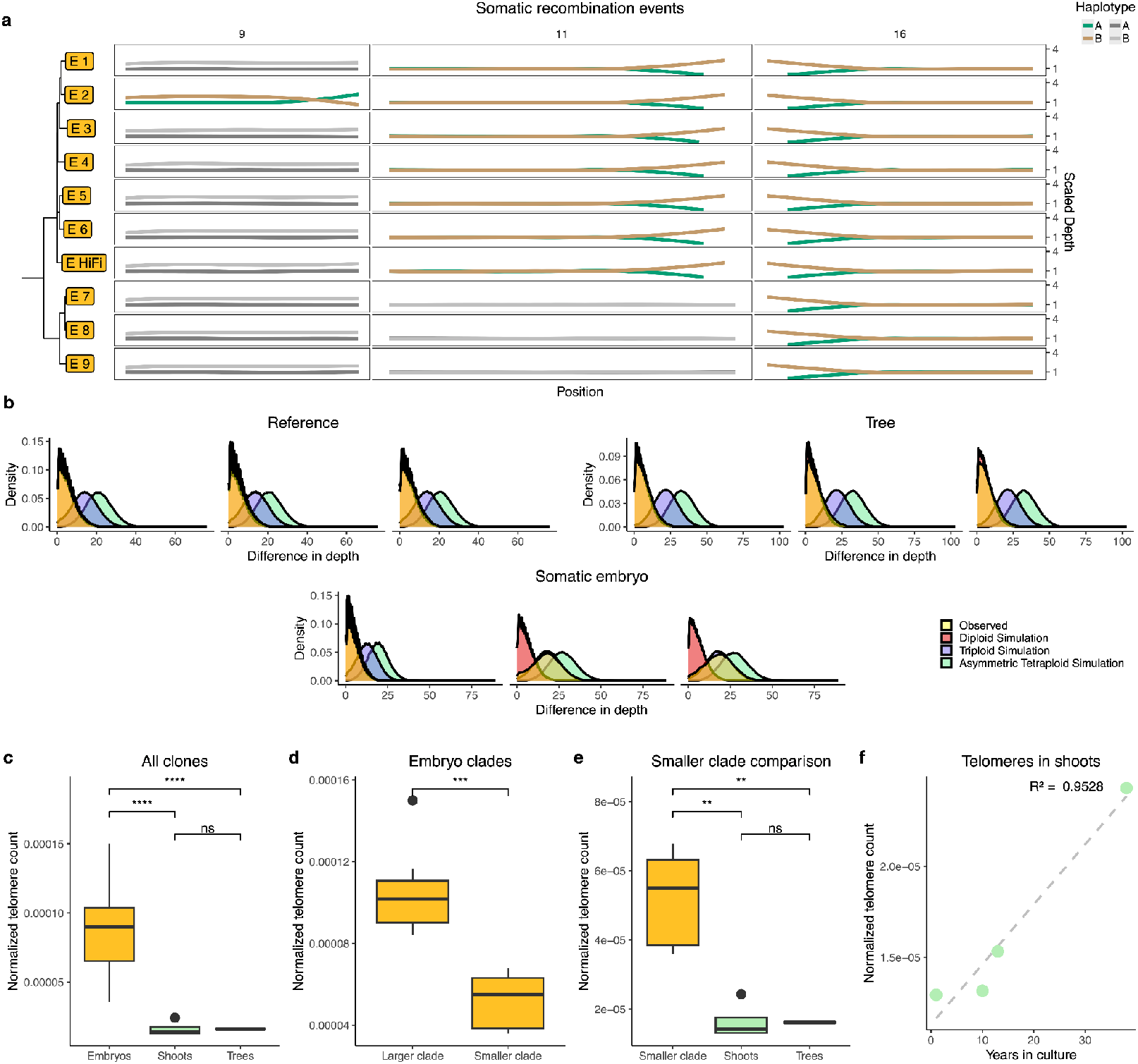
Somatic recombination, ploidy simulations, and telomeres. **a**, Haplotype-resolved depth across chromosomes exhibiting somatic recombination in the somatic embryos. **b**, For all long-read sequenced samples, the difference between the reference site depth and alternate site depth was taken. The expected differences in these depths were calculated based on proposed diploid (equal weight), triploid (1/3 and 2/3), and asymmetric tetraploid (1/4 and 3/4) ratios sampled from a binomial distribution. These simulated distributions were plotted along with the distributions of the observed data in the three long-read sequenced samples. The chromosomes duplicated in the somatic embryos are shown in the somatic embryo and tree clones, as well as chromosome one to represent a typical diploid chromosome **c**, Number of telomeric repeats pooled by method of clonal propagation. A Welch’s t-test was performed comparing the somatic embryos to the shoots and trees. ****: p <= 0.0001, ***: p <= 0.001, **:p <= 0.01, *: p <= 0 05. **d**, Somatic embryos were separated into the two largest clades. The telomeric repeats in each clade were pooled and compared to one another. A Welch’s t-test was performed comparing the two clades. **e**, The telomere repeats of the smaller of the two somatic embryo clades were pooled and compared to the shoots and trees. **f**, The telomeric repeats in the shoot clones. The relationship between years in culture and telomeric repeal count was assessed with a linear model, and model fit was determined.

**Supplementary Fig. 11.**
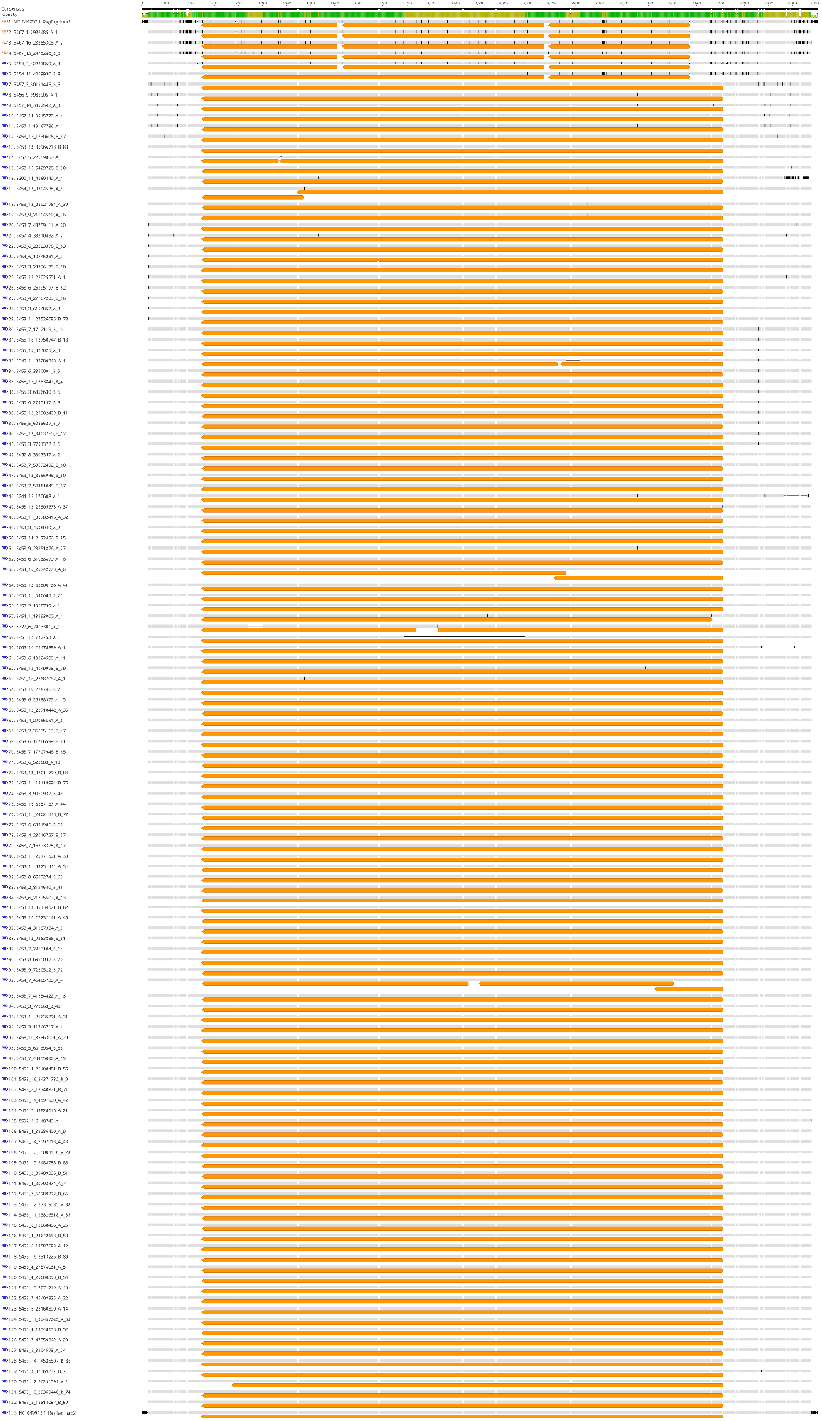
Distinct open reading frame structures exist in the 5500 class TE. Alignment of all trimmed 5500 class TEs and the matching sequences from the reference genome assembly visualized with Geneious. Open reading frames are annotated in orange boxes underneath the sequences. The sequences matching consensus are represented in grey, with alternate bases displayed in black.

**Supplementary Fig. 12.**
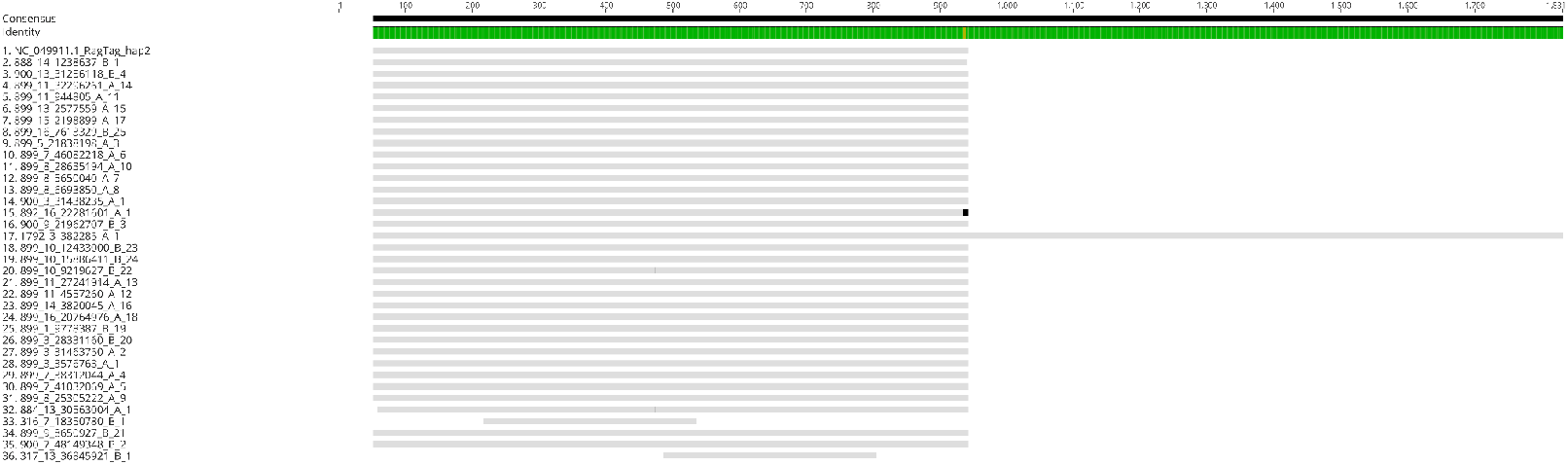
No observable open reading frame structures in the 900 class TE. Alignment of all trimmed 900 class TEs and the matching sequence from the reference genome assembly visualized with Geneious. No open reading frames were observed, thus there is no annotation. The sequences matching consensus are represented in grey, with alternate bases displayed in black.

**Supplementary Fig. 13.**
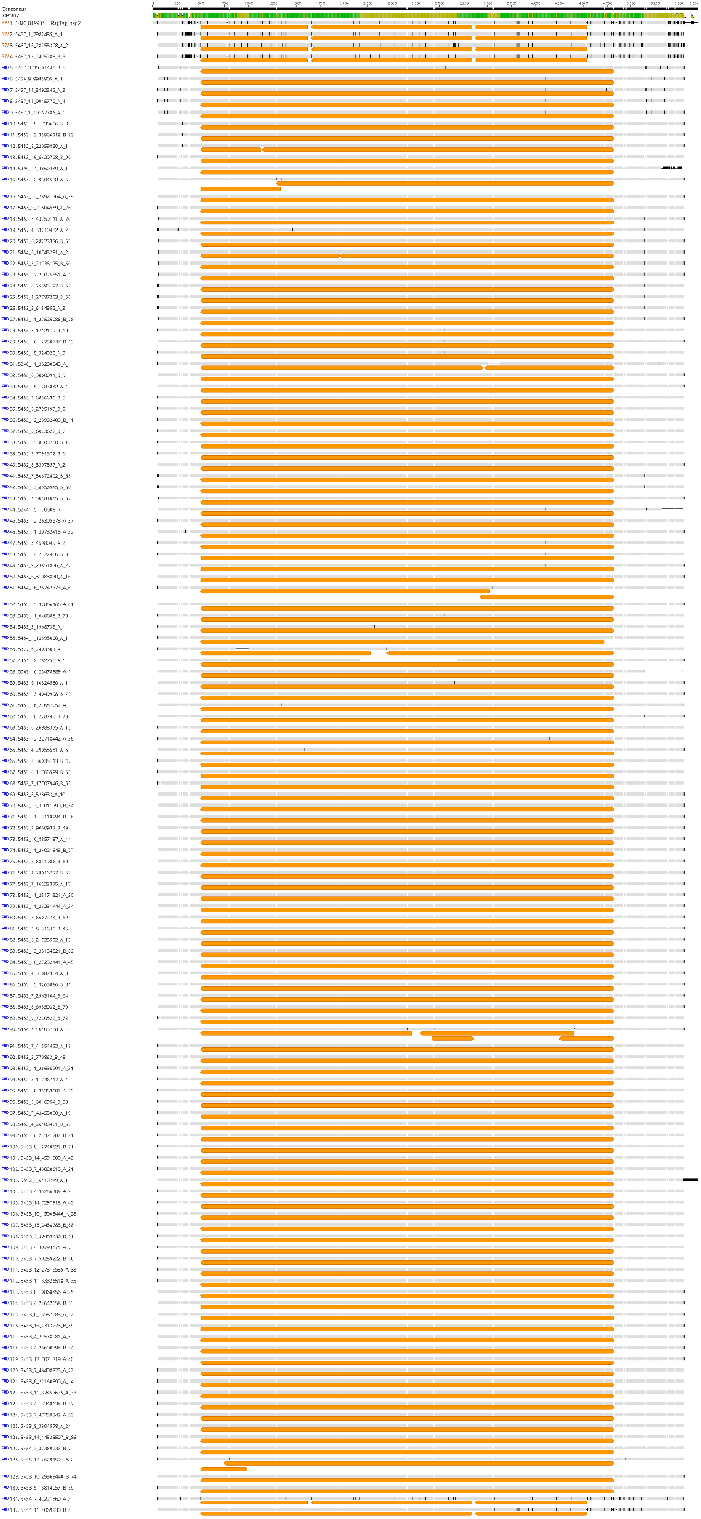
Untrimmed class 5500 insertions show target site duplications. Alignment of all untrimmed 5500 class structural variants and the matching sequences from the genome assembly visualized with Geneious. Open reading frames are annotated in orange boxes underneath the sequences. The sequences matching consensus are represented in grey, with alternate bases displayed in black.

**Supplementary Fig. 14.**
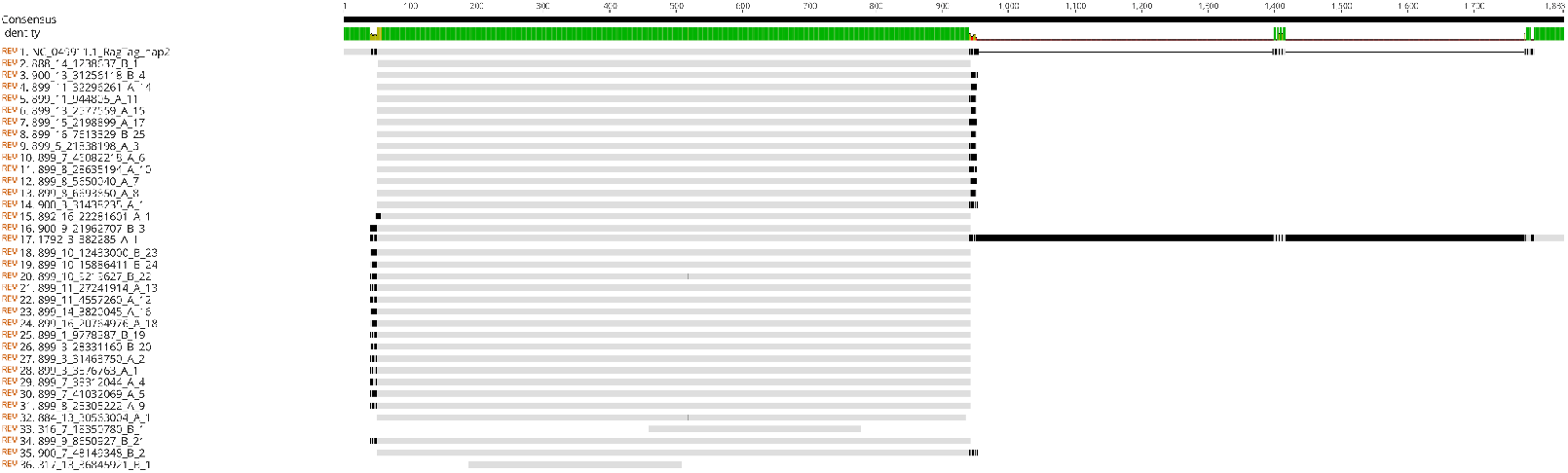
Untrimmed class 900 insertions show target site duplications. Alignment of all untrimmed 900 class structural variants and the matching sequence from the genome assembly visualized with Geneious. The sequences matching consensus are represented in grey, with alternate bases displayed in black.

**Supplementary Fig. 15.**
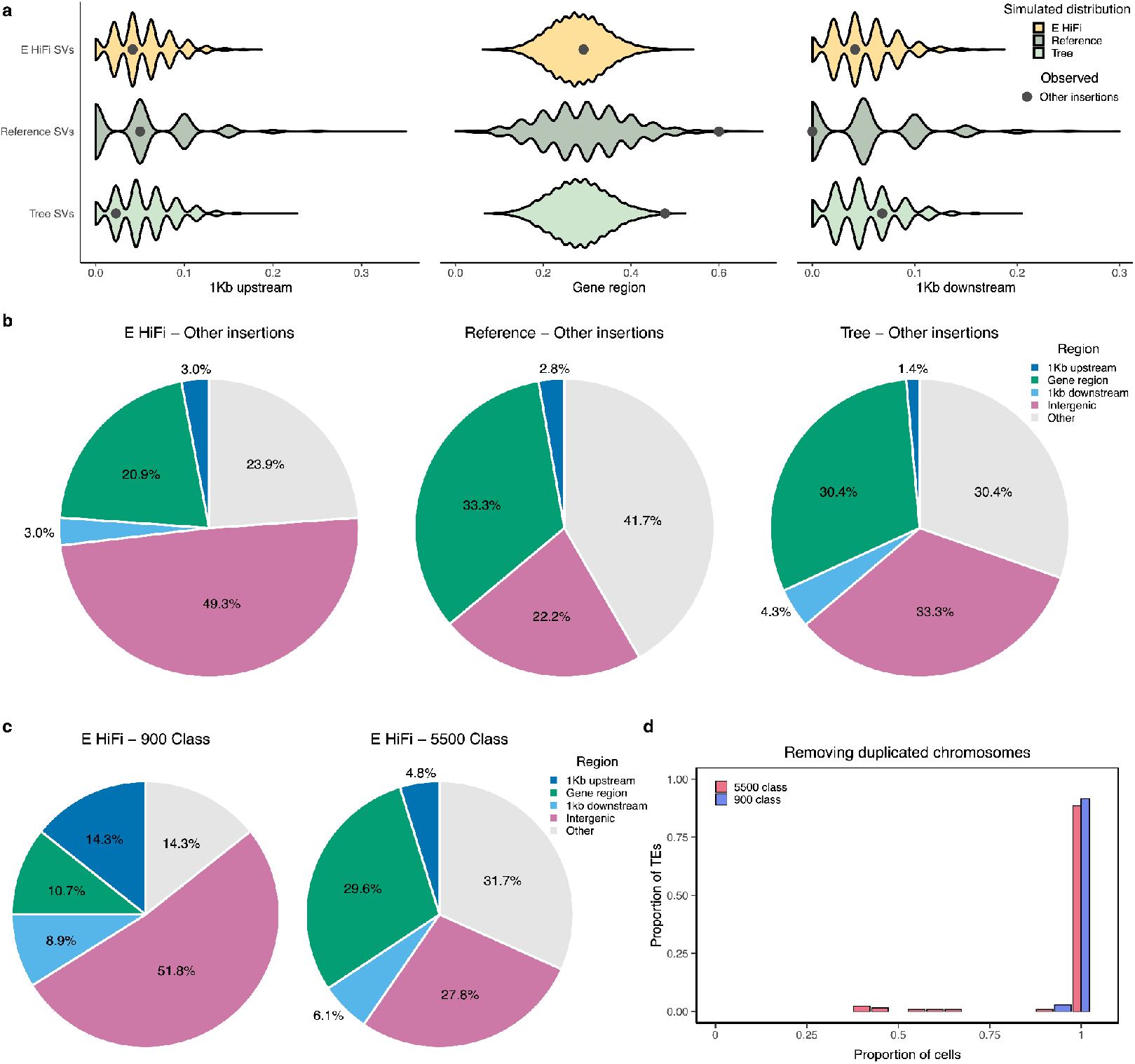
Transposons exhibit different biases compared to other insertions. **a**, *De novo* structural variant insertions in the long-read sequenced samples not classified as transposons were simulated to randomly insert across the genome 10,000 times, shown by the violin plots. The points indicate the observed means of the insertions. **b**, The proportion of the *de novo* structural variant insertions not classified as transposable elements in various genomic features in the long-read sequenced samples. **c**, The proportion of the 900 and 5500 class insertions in various genomic features in the long-read sequenced embryo. **d**, The 900 and 5500 class TEs observed in different proportions of cells that compose the somatic embryo with the duplicated chromosomes removed.

**Supplementary Fig. 16.**
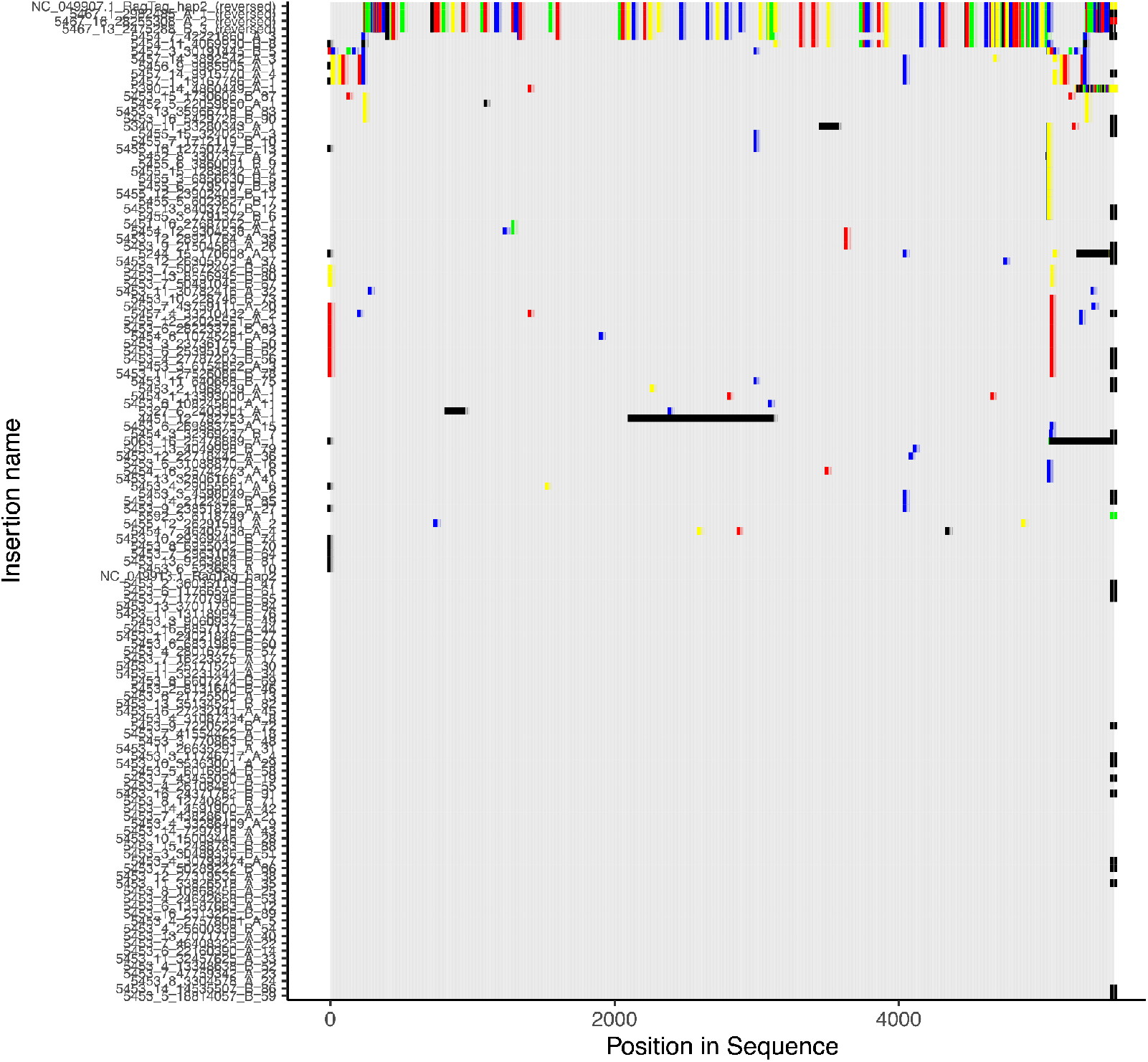
Observable SBS and InDels in the 5500 class TEs. Alignment of all trimmed 5500 class TEs and the matching sequences from the genome assembly. Consensus was determined as the most common base at that position and visualized as grey. Deletions were visualized as black Differences from the consensus of A were plotted in red, T were plotted in blue, C were plotted in green, and G were plotted in yellow. For visualization purposes, any variation from the consensus was plotted more thickly than consensus sequences.

**Supplementary Fig. 17.**
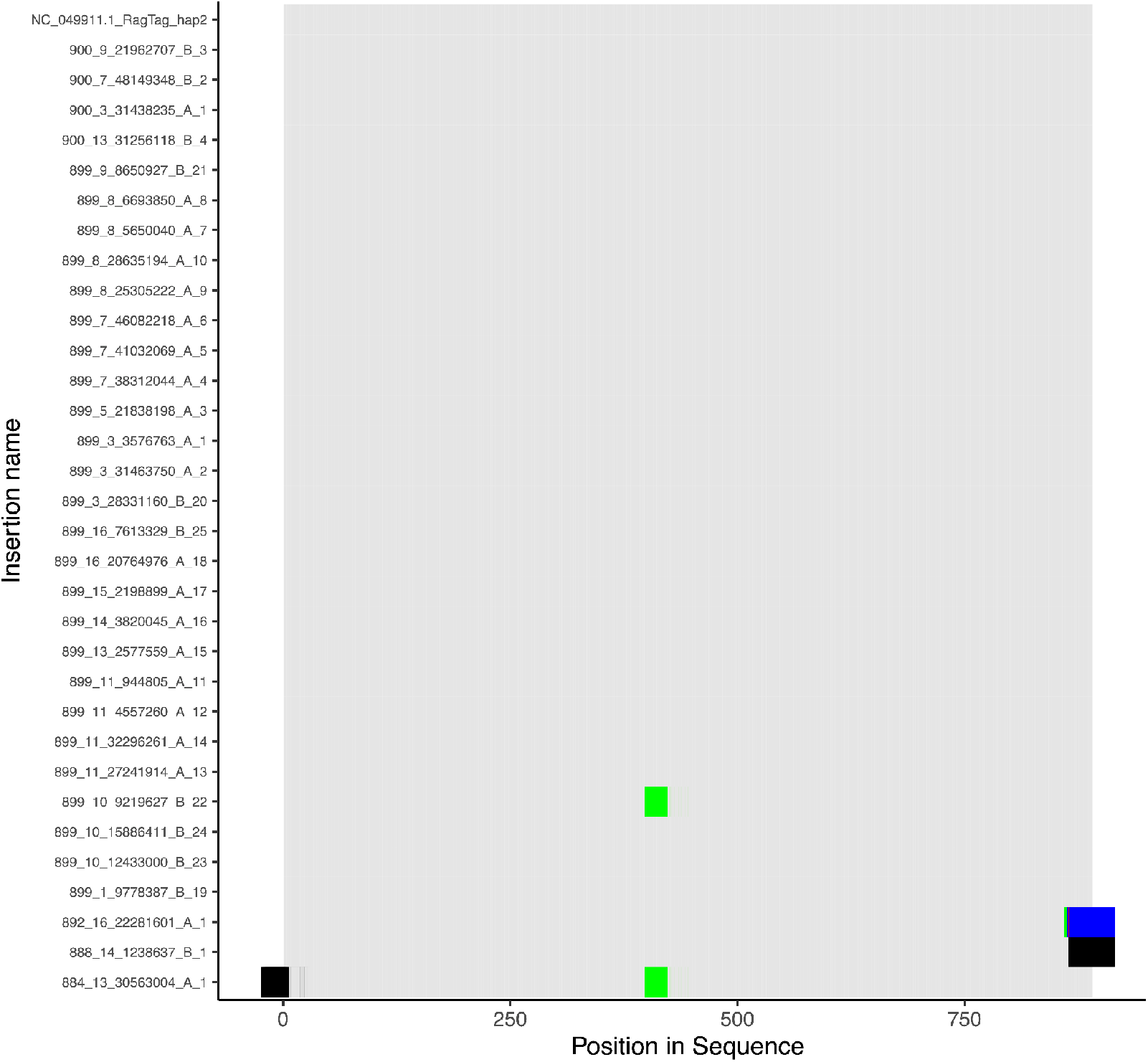
Observable SBS and InDels in the 900 class TEs. Alignment of all trimmed 900 class TEs and the matching sequences from the genome assembly. Consensus was determined as the most common base at that position and visualized as grey. Deletions were visualized as black Differences from the consensus of A were plotted in red, T were plotted in blue, C were plotted in green, and G were plotted in yellow. For visualization purposes, any variation from the consensus was plotted more thickly than consensus sequences.

**Supplementary Fig. 18.**
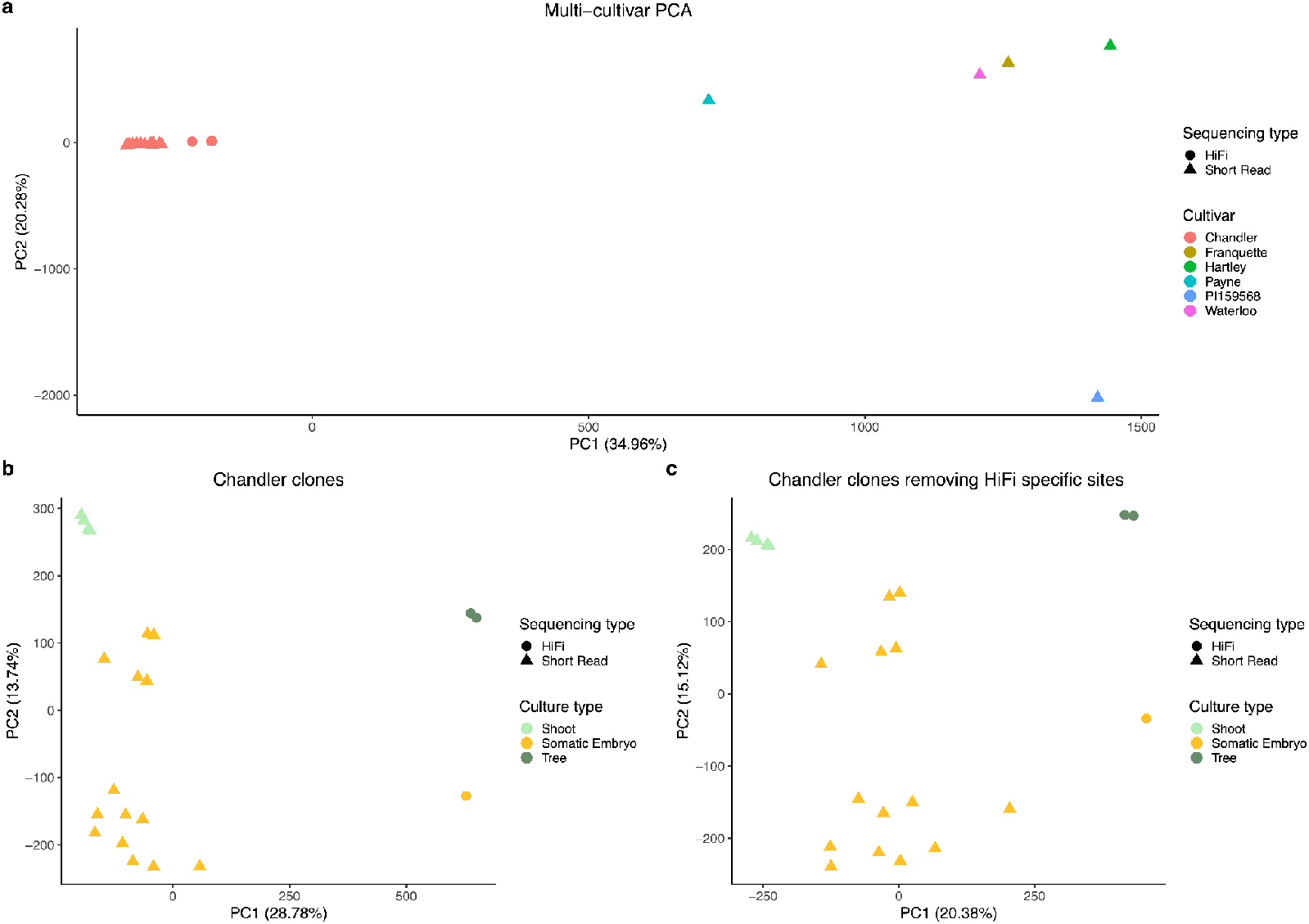
PCA shows the clones are *Juglans regia* ‘Chandler’ and sequencing effects. **a**, PCA of multiple walnut cultivars and all clones. Sequencing method is denoted by shape while cultivar is denoted by color. **b**, PCA of all clones, with shape denoting sequencing method and color denoting clonal propagation method. **c**, PCA of all clones with sites that were called in all long-read sequenced samples and no short-read sequenced samples removed. Shape denotes sequencing method and color denotes clonal propagation method.

### Supplementary Tables

Supplementary Table 1 - Sequencing statistics of all walnut clones and sequencing methods

Supplementary Table 2 - Statistics of primary and haplotype-resolved Juglans regia ‘Chandler’ assemblies

Supplementary Table 3 - The genetic map of Juglans regia ‘Chandler’ constructed from self-pollinated progeny

Supplementary Table 4 - The percent of sites properly phased in the reference genome as determined by the haplotype assignment in the genetic map

Supplementary Table 5 - De novo mutation counts of SBS, InDels and SVs with the filtering thresholds median quality >= 30, median depth >=15 and 100% mappable regions with 150bp reads

Supplementary Table 6 - Retained de novo mutations on chromosome 4 occurring after duplication but still present in the diploid somatic embryos

Supplementary Table 7 - Number of de novo mutations on chromosomes 4 and 9 occurring before chromosome duplication

Supplementary Table 8 - Counts of 21-mer telomeric repeats from nuclear genome aligned reads in the walnut clones

Supplementary Table 9 - Names and labels found in scripts and in repositories and their correspondence with the manuscript

